# tinyRNA-seq: An optimized approach to sequencing tiny RNAs and primitive RNA genomes

**DOI:** 10.64898/2026.08.06.743385

**Authors:** Ben W.F. Colville, Jamie Zhao, Liam Hade, Jack W. Szostak

**Affiliations:** HHMI, The University of Chicago, Chicago, IL 60637; Department of Chemistry, The University of Chicago, Chicago, IL 60637; Institute for Biophysical Dynamics, The University of Chicago, Chicago, IL 60637

## Abstract

Very short RNAs play critical roles in modern biology, and are thought to have been crucial for genome replication during the origin of life. Next-generation sequencing is an essential tool for characterizing pools of small RNAs, but current library preparation methods suffer from strong size and sequence biases. Here we present tinyRNA-seq, an optimized library preparation method designed to minimize length- and sequence-dependent capture bias enabling the sequencing of RNA fragments as short as 2 nucleotides. We use degenerate adaptor regions to reduce ligation sequence bias and facilitate unique molecular identifier (UMI) installation. We benchmarked tinyRNA-seq against commercial kits using a model primordial RNA genome consisting of hundreds of defined oligonucleotides ranging from 2 to 12 nucleotides. tinyRNA-seq reproduced the input RNA distribution without the size and sequence bias of the commercial kits. tinyRNA-seq also enables the detection of *de novo* oligonucleotide generation, an important process for the origins of life. Applied to biologically derived small RNAs including miRNAs, piRNAs, and cityRNAs, tinyRNA-seq showed significantly lower capture bias and recovered a wider range of sequences than commercial kits. tinyRNA-seq may thus provide a more complete and quantitatively accurate representation of small RNAs from both biological and chemical sources.

**GRAPHICAL ABSTRACT:** 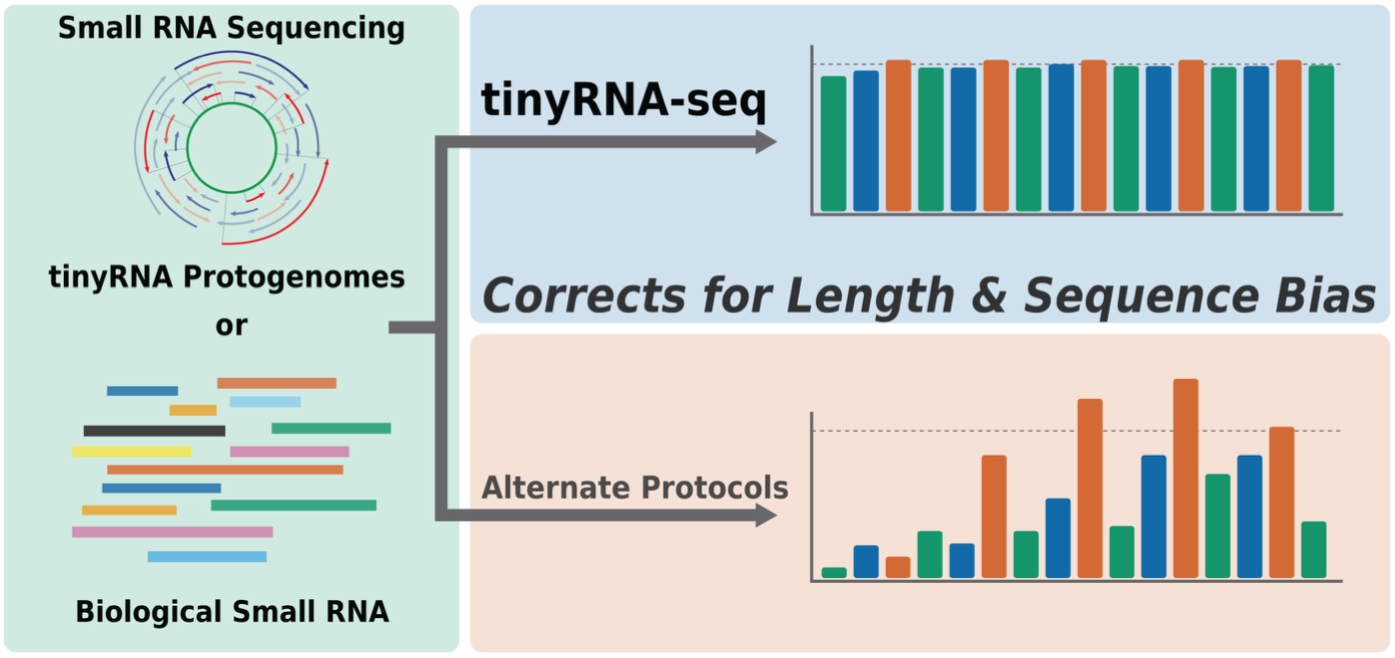

## INTRODUCTION

The RNA world hypothesis suggests that the earliest forms of life would have used RNA as both an informational polymer and for primitive catalytic functions[1]. Early RNA genomes were likely not one specific sequence or length but rather a complex mixture of species whose emergent properties would have determined their viability. RNA protogenomes contained within protocells would likely have been short in length, with the upper limit of genome size set by RNA copying efficiency, fidelity and genomic stability [2, 3].

The absence of protein enzymes and evolved ribozymes during this early stage of life implies that template-dependent chemical copying of RNA must have been responsible for replicating early RNA genomes[4]. Non-enzymatic RNA primer extension (NERPE) has been proposed as a model of primitive genome copying via the templated polymerization of prebiotically plausible imidazole-activated nucleotides. The discovery of the key reactive nucleotide intermediate in this process, 5′-5′-phosphorimidazolium-bridged-dinucleotides (N*N), has led to greater understanding of the mechanism of NERPE[5–9]. Despite this progress, there remain several problems with NERPE as a means of copying long RNAs, including: strand separation, the ‘last base addition problem’[10] and *de novo* oligonucleotide generation[11].

When copying longer, ribozyme-length RNA, strand separation becomes problematic as high duplex melting temperatures render long sequences impossible to copy, with strands re-annealing on timescales orders of magnitudes faster than copying chemistry. Environmental fluctuations in pH[12], solvent viscosity[13] or salinity[14] as well as the incorporation of a small amount of non-canonical 2′,5′-linkages in the RNA backbone[15] may allow some enhanced strand separation and therefore copying of longer RNA, but the problem remains fundamentally unsolved in the context of the cycles of copying required for replication.

The ‘last base addition problem’ suggests that information loss from the 3′-terminus of RNA due to slower copying of terminal positions relative to internal sites could prevent complete copying of functional length RNA over many cycles. This finding is consistent with the mechanism for NERPE which proceeds via imidazolium-bridged dinucleotides (N*N) that bind the template with two base pairs[16, 6]. As primer extension via activated monomers (*N) leads to slow and low fidelity copying it is highly desirable for primer extension to proceed via the binding of bridged dinucleotides. A system that avoids this last base problem would be advantageous in maintaining the information of a protogenome[10, 17].

Genome maintenance at the 5′-end seems even more challenging as it requires the continuous supply of a specific primer sequence that can outcompete primer extension initiated at internally primed sites. Viroid-like rolling circle copying has been explored as a potential solution but would rely on extensive primer extension of both strands and tolerance of the topological challenges of small circular RNA genomes[18, 19].

For any RNA replication system to sustain itself it must be supplied with sufficient new primers to double the genetic information for every cycle of replication. While many studies have shown that untemplated polymerization of RNA could generate short random primers it would be highly advantageous if this process were compatible with NERPE[20–23]. In particular, template-dependent rather than random *de novo* oligonucleotide generation would avoid the introduction of exogenous sequences into an early genome[24, 25].

In an attempt to satisfy these requirements of primitive RNA genome replication we have recently developed a theoretical framework to solve the multiple problems with non-enzymatic templated copying of RNA, dubbed the “Virtual Circular Genome” (VCG) model[26]. Here, the genome is distributed over many short oligos from both strands of a virtual circular sequence (**Figure 1A**). As the genome is circular, copying can be initiated at any point. Further, extending each oligo in the pool by only one nucleotide, coupled with *de novo* oligonucleotide generation to feed the system, can in principle replicate the genome. We envisage that seeding of protocells with VCG-like genomes might occur by the encapsulation of random sequence oligonucleotides generated by untemplated mechanisms such as eutectic phase polymerization[27, 28].

**Figure 1:**
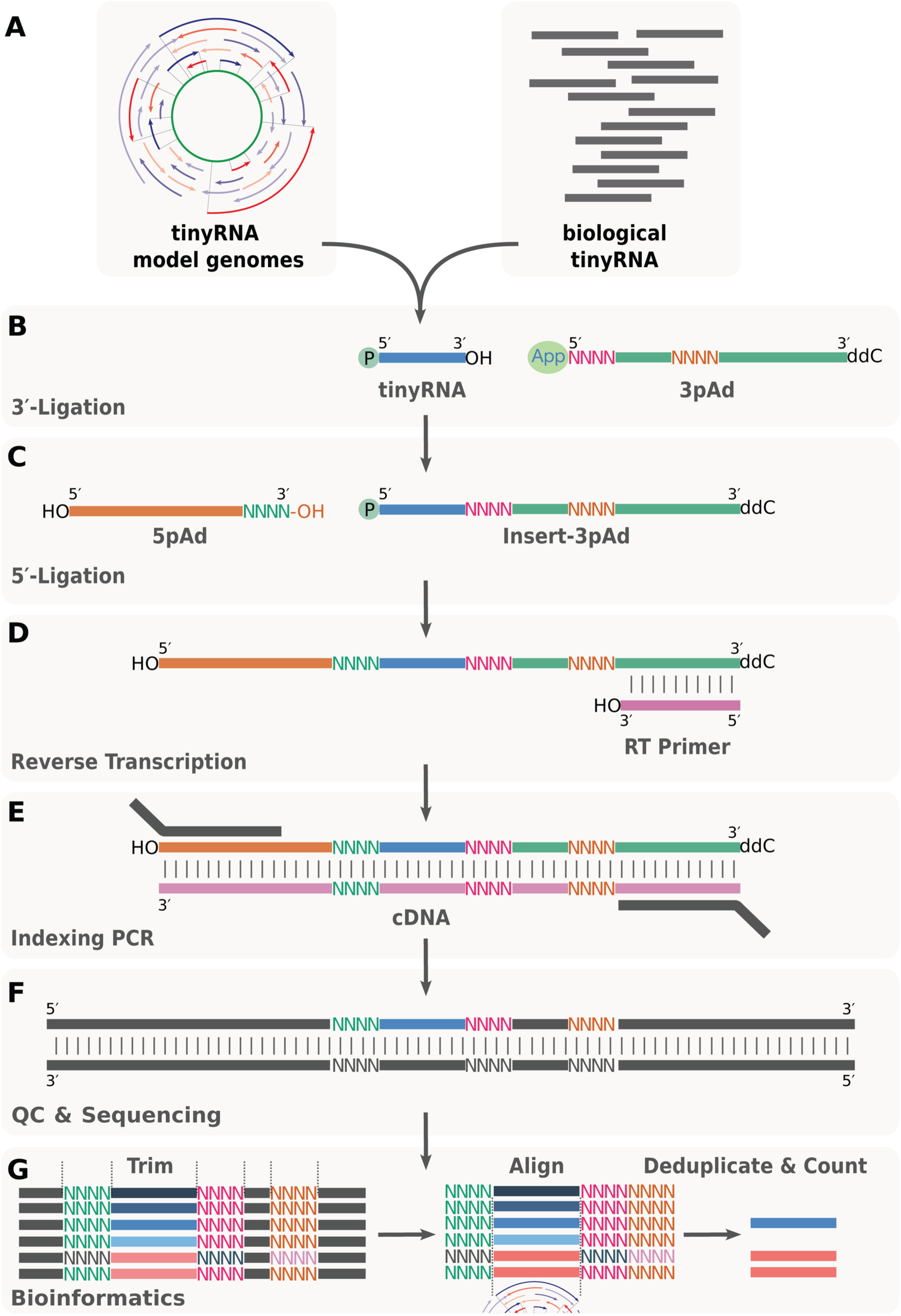
Protocol for preparing sequencing libraries using tinyRNA-seq. (**A**) small RNAs with 5′-phosphate and 3′-hydroxyl termini such as components of primitive RNA genomes or small RNAs from biological sources are purified. (**B**) In the first step input RNA undergoes enhanced enzymatic ligation to the 3′-adaptor (**3pAd**), a pre-adenylated ssDNA with a randomized 5′ end, using a mutant T4 RNA ligase 2 (truncated, KQ). (**C**) Excess adaptor is removed using a combination of enzymatic de-adenylation (5′-Deadenylase) and DNA 5′-end specific exonuclease (RecJf) and the RNA-DNA hybrid is ligated to the 5′-adaptor (**5pAd**), an RNA with a randomized 3′-terminus, using T4 RNA ligase 1. (**D**) The **3pAd** contains the reverse transcription primer (RT Primer) binding site. Reverse transcription followed by RNA destruction allows recovery of purified cDNA. (**E**) Real-Time PCR is used to amplify the cDNA to the desired amount without introducing amplification bias while simultaneously adding flowcell binding sequences for Illumina sequencing. (**F**) Purified dsDNA is quality controlled, pooled and sequenced on the Illumina sequencing platform. (**G**) A custom bioinformatics pipeline allows handling of degenerate regions and reassembling a unique molecular identifier (UMI) for each read which can be used to collapse PCR duplicates.

Experimental tests with a synthetic VCG pool, as well as computational simulations show that primer extension is possible in a model VCG system. Furthermore, the collective encoding of information in a VCG could provide greater resilience to copying errors[29, 30]. To further explore the dynamics of VCG genomes, we now require a more complete experimental picture. Indeed, small changes in sequence distribution could have important ramifications for overall genome function and information storage. We have previously studied NERPE copying using deep sequencing of self-priming RNA hairpins (NERPE-seq) but this cannot be used to investigate a complex and chemically evolving mixture of RNAs such as a VCG[17, 31–33]. Therefore, a new sequencing methodology capable of capturing very short RNA (tiny RNA) with high accuracy and low bias is essential for exploring the dynamics of RNA protogenomes.

Small RNA sequencing is a method used for studying biological small RNAs (sRNAs) using next generation sequencing[34]. These RNAs, usually <35 nt in length, such as microRNA (miRNA), PIWI-interacting RNA (piRNA) and other emerging types such as cleavage-inducing tiny guide RNAs (cityRNAs) represent a class of noncoding RNAs critical to eukaryotic gene regulation[35–39]. The presence and abundance of these species are also of great interest as biomarkers in disease pathology and have prompted the development of many different approaches to their sequencing[40–43]. All small RNA sequencing involves the installation of known sequences on the 5′- and 3′-ends of the RNA of interest, typically by stepwise enzymatic ligations (**Figure 1A-B**). This enables reverse transcription to cDNA (**Figure 1C**), amplification by PCR, multiplexing and flowcell binding for sequencing (**Figure 1D**). Despite these relatively simple steps and many efforts to optimize sequencing of small RNAs it remains challenging to accurately capture species of this short length scale[44–47].

Here we present an optimized small RNA sequencing protocol to study an ensemble of VCG tiny RNAs with as close to quantitative accuracy as possible. We introduce tinyRNA-seq, a method for the unbiased capture and deep sequencing of tiny RNAs (<17 nucleotides). We benchmark tinyRNA-seq on our model VCG system, a pool of 247 oligos with lengths between 2 and 12 nucleotides, as well as on biologically derived small RNAs from mouse brain tissue. We identify and address known sources of bias in small RNA sequencing to ensure that sequencing read counts closely reflect the actual abundances of oligonucleotides in the RNA pool. We develop a ligation-barcoding method that simultaneously mitigates ligation capture bias while allowing correction for duplicate sequencing reads. We demonstrate the power of real-time PCR amplification to prevent the introduction of bias for small RNAs of interest. To enable this, we have also created a data analysis module that streamlines analysis of multiplexed samples. We show that tinyRNA-seq can be used to analyze previously intractable problems in origins of life studies, such as the *de novo* oligonucleotide generation that is essential for sustaining VCG RNA genomes. We discuss the uses of tinyRNA-seq in the context of both studying the origins of life and small RNAs in general.

## MATERIAL AND METHODS

### The VCG pool and tinyRNA-seq pipeline overview

We are interested in developing an unbiased method for the deep sequencing of tiny RNAs (<17 nt), for example, components of the Virtual Circular Genome (VCG)[26, 29]. We selected the same model VCG genome sequence we have investigated previously, which consists of a pool of 247 synthetic tiny RNAs with 5′-phosphate and 3′-hydroxyl ends that all map to either the sense, antisense or both strands of the 12 nt circular genome (**Table S4**)[29]. The oligos range in length from 2 to 12 nucleotides with oligos of length 2 having 11 distinct sequences, length 3 having 20, and lengths 4 through 12 comprising 24 sequences, with 12 from the sense and 12 from the antisense strand. Using this synthetic pool of equimolar tiny RNAs allows for a rigorous investigation of the capture bias that is known to be problematic for this RNAs of this length regime.

### 3′-end ligation

2 µL of 10 µM VCG RNA was mixed with 2 µL 10x T4 RNA Ligase buffer (NEB B0216SVIAL), 7 µL 50% PEG8000 (NEB B1004SVIAL) and 2 µL T4 RNA Ligase 2 truncated KQ (NEB M0373S) and 3′- end adaptor (**Table S3**, we used a 1:1 mixture of both **3pAd-CAA** and **3pAd-GTT**) to a final concentration of 2 µM and made up to a final volume of 20 µL. The reaction mixture was incubated at 16 °C for 16 h and then 65 °C for 5 min to denature the ligase.

### 3′-end ligation adaptor depletion

To remove excess **3pAd** adaptor we use a combination of a de-adenylating enzyme followed by a 5′ to 3′ ssDNA specific exonuclease using the following protocol. To each ligation reaction was added 1 µL of 5′ Deadenylase (NEB M0331S) and incubated at 30 °C for 1 h. Then 1 µL of 1.25 M NaCl and 1 µL of RecJf (NEB M0264S) was added and incubated for 1 h at 37 °C. The mixture was purified using the modified spin protocol outlined below and eluted in 10 µL water.

### Modified spin column protocol

The reactions were then purified with Oligo Clean and Concentrate kits (Zymo Research) with minor modifications to the manufacturer’s protocol, outlined herein. Each reaction was diluted to 50 µL with water and 200 µL Oligo Binding Buffer (Zymo Research) was added. This was then combined with 400 µL 100% ethanol and loaded onto an IC-Spin column (Zymo Research). The tube was then either a) loaded into the receiver tube and centrifuged at 12,000 rcf for 1 min or b) connected to a vacuum manifold for more facile parallel processing of multiple samples. For both a) and b) the columns were washed with 800 µL DNA Wash Buffer (Zymo research) and spun (12,000 rcf for 1 min) or vacuum filtered again. For both methods the resulting tubes were transferred to new receiver tubes and spun again at 12,000 rcf for 2.5 mins. RNA or cDNA was eluted from the now dry columns into a new 1.5 mL Eppendorf tube by adding the desired volume of water plus and additional 1 µL of water as the column absorbs and retains ∼ 1 µL of applied water in our hands. Once the water was added directly to the column, the tubes were incubated at RT for 1min before spinning at 12,000 rcf for 1 min. The RNA or cDNA was either used immediately or frozen at −30 °C for future use.

Normal elution volumes for the clean-up of respective steps are: 3′-end ligation; 10 µL water, 5′- end ligation; 20 µL water, RT; 16 µL water.

### 5′-end ligation

The 10 µL of purified RNA-DNA hybrid from the 3′-end ligation was transferred to a PCR tube and the following added: 5 µL 25 µM 5′-end adaptor (**5pAd**, **Table S3**), 3 µL 10x T4 RNA Ligase buffer (NEB B0216SVIAL), 5 µL 50% PEG8000 (NEB B1004SVIAL) and 1 µL T4 RNA Ligase 1 (NEB M0204S), 1 µL 10 mM ATP (NEB B0756AVIAL) to 25 µL final volume. The mixture was incubated at 16 °C for 16 hours and then 65 °C for 5 mins to denature the ligase. The mixture was purified using the modified spin protocol outlined above and eluted in 20 µL water.

### Reverse transcription

To the 20 µL of purified 5′-end ligation in a PCR tube was added 0.7 µL of 100 µM reverse transcription primer (RT_Primer, **Table S3**). The RT primer was annealed with the following program: 75 °C for 3 mins, 37 °C for 10 mins, 25 °C for 10 mins. To this, 2.5 µL 10x DTT (NEB B1034AVIAL), 5 µL 5x ProtoScript^®^ II buffer (NEB B0368SVIAL), 1 µL 10 mM dNTPs (NEB N0447S) and 2 µL ProtoScript^®^ II (NEB M0368S) were added and the mixture incubated at 42 °C for 12 h and then 65 °C for 5 mins to denature the enzyme. The mixture was purified using the modified spin protocol outlined below including the RNA destruction step (10 µL 0.5 M EDTA, 10 µL 1 M NaOH, 65 °C 15 mins) and eluted in 10 µL water.

### Amplification by PCR

PCR was performed using a Thermo Scientific QuantStudio Pro 7 qPCR machine enabling real time monitoring in order to prevent over-amplification, a well-known contributor to PCR bias. With RNA input of 20 pmol we were able to only run 8-12 cycles. Additionally, in our hands we found that a short incubation with RNase H improved the quality of the libraries generated.

To 4 µL of purified cDNA was added a PCR mixture containing 1.25 µL of 10 µM of the forward and reverse primers (NEBNext Multiplex Oligos for Illumina Sets 1 & 2), 0.5 µL RNase H (ThermoFisher), 5 µL 5x Q5 reaction buffer (NEB B9027SVIAL), 1 µL 10 mM dNTPs (NEB N0447S), 1.25 µL EvaGreen (Biotium, #31000), 0.5 µL Q5 HiFi Hot Start Polymerase (NEB M0493S) and 10.25 µL water to a total volume of 25 µL.

Samples were first incubated with RNase H (Thermo Fisher) at 37 °C for 5 minutes, followed by denaturation at 98 °C for 30 seconds. PCR amplification was then performed over a max of 16 cycles consisting of denaturation at 98 °C for 10 seconds, annealing and extension at 72 °C for 40 seconds. A final extension step was carried out at 72 °C for 2 minutes before holding at 4 °C.

5 µL of Purple Loading Dye (NEB B7024S) was added to the sample and the entire volume was loaded and run on a 3% TBE agarose gel (2:1 MetaPhor^®^ Agarose Lonza #50181: UltraPure™ Agarose, Invitrogen, #16500500) for 1-1h 30 mins at 80 V. The relevant band was visualized under blue light, excised and chopped to fine pieces with a razor blade then loaded into Freeze ′N Squeeze™ DNA Gel Extraction Spin Columns (BioRad #7326165). The tube was placed on dry ice for 30 minutes and then the dsDNA recovered by spinning (13,000 rcf for 3 mins, RT) as per the manufacturers protocol.

### Library quantification and sequencing

Purified PCR products were quantified using an Agilent TapeStation 4200 using the D1000 reagents (Agilent, ScreenTape: 5067-5582, Reagents: 5067-5583) and sequenced on an Illumina MiSeq using a 150-cycle paired end kit v3 (Illumina MS-102-3001).

### Bioinformatics

#### CPM and residual

To normalize between samples with different total sequencing reads we use Counts per Million (CPM) defined as:

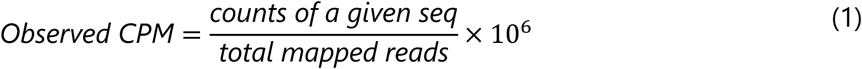

Due to our known input pool (247 known VCG sequences) we can also naively calculate expected CPM for each VCG sequence as:

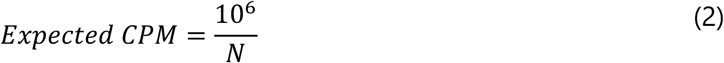

where N is the number of unique sequences. However, as we have 2 and 3 nt sequences that appear multiple times (copy number > 1) and therefore have a commensurately higher input concentration, we define Expected CPM more accurately as:

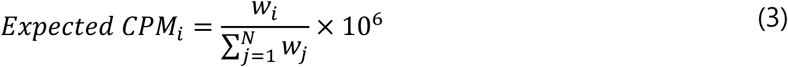

where ***w****_i_* is the copy number multiplier for a sequence *i* and the denominator is the sum of all weights across all N pool members.

We can then calculate the residual for each VCG oligo, which acts as a simple metric for bias, where:

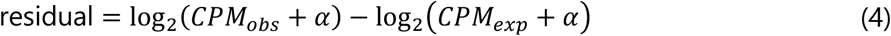

Where CPM_exp reflects the correctly weighted input ratios and *α* is a pseudocount to handle sequences with 0 reads that are retained in our analysis. A residual of 0 reflects perfect representation with respect to expected, residual values greater than 0 represent VCG sequences that are overrepresented compared to expected and residual values lower than 0 represent sequences that are underrepresented. Known sequences not captured receive a large negative residual proportional to their expected CPM rather than excluding them from analysis. Here we opted to use *α* = 1 making the expected CPM for a VCG sequence present once ≈ 4048 per million reads.

## RESULTS

### tinyRNA-seq enables accurate compositional profiling of short synthetic RNA pools

### Overview of the tinyRNA-seq protocol

To address the key sources of bias in current small RNA sequencing protocols we set out to design adaptor combinations and a workflow that would solve multiple problems simultaneously. We focused on: (1) reducing primary sequence bias during ligations, (2) reducing secondary structure bias during ligations[45, 46], (3) reducing input RNA length bias during ligations, (4) correcting for read duplication introduced by PCR[40, 48], and (5) maintaining forwards compatibility with future experiments. It is well established that T4 RNA ligase enzymes have sequence preferences at ligation junctions. A common approach to address this is to use adaptors with a degenerate region (typically 3-7 nt) so that more potentially favourable ligation junctions can be sampled during the critical first ligation step (**Figure 1B**)[40, 47, 41, 43]. Inspired by this, we designed a custom 3′-dideoxy and 5′-adenylated ssDNA adaptor, **3pAd** (**Supplementary Information, Table S3**): the **3pAd** contains two degenerate regions of 4 nucleotides (one at the 5′-end and one embedded internally), as well as a constant region for bioinformatic handling, a 3-nucleotide multiplexing barcode, and the reverse transcription primer binding site. We anticipated that the degenerate 5′-end in concert with the commonly used mutant of T4 RNA ligase 2 (T4 Rnl2, truncated KQ) would reduce sequence bias, and that the 3′-dideoxy end would simultaneously prevent self-ligation of the **3pAd** adaptor.

To address the problem of secondary structure bias for small RNA-seq, we sought to take advantage of work elucidating the natural substrate of T4 RNA ligase 1: tRNA hairpin loops[49]. We predicted we could increase the frequency of favourable co-folding events by using a 5′-adaptor that also ended in 4 degenerate bases (**5pAd, Supplementary Information, Table S3**) as this, in combination with the embedded internal degenerate region of **3pAd**, could allow more “hairpin-like” structures to form during the second ligation step (**Figure 1C**). We also placed the 4N region of **5pAd** downstream of the Illumina sequencing primer binding site to enable recovery of all 12 degenerate bases (**5pAd**: 4, **3pAd**: 4+4) in our final sequencing construct. Using this we implemented a bioinformatic step to eliminate PCR duplication events known to be problematic for small RNA inserts (**Figure 1F**). Additionally, the first 4 bases of Illumina sequencing are known to be particularly sensitive to low sequence diversity, which our 4N region of **5pAd** addresses. Lastly, we implemented longer incubations times at lower temperatures (16 h at 16 °C vs 1 h at 25 °C for typical protocols) with higher amounts of enzyme for our ligations to help capture our very short RNAs of interest.

#### Benchmarking tinyRNA-seq against commercial kits reveals correction of length and sequence-dependent capture bias

We noted that all small RNA-seq studies on identifying bias as well as all currently commercially available kits focus on the length scale of miRNAs (21-23 nt). As our particular interest is with very short, prebiotically plausible oligos (2-12 nt) we tested tinyRNA-seq against a control protocol using adaptors with defined termini (we dub “DEF-seq”) and commercial kits to examine if the same biases were relevant to the sequencing of our VCG system. The control protocol (DEF-seq, **Supplementary Information**) we used is derived from the small RNA sequencing literature and is similar to tinyRNA-seq in that is ligation based, it differs in that: it uses adaptors with no degenerate regions, unoptimized ligations and no real-time PCR [50]. We prepared sequencing libraries using either tinyRNA-seq, DEF-seq or one of 5 commercial kits: 4 ligation-based kits from different manufacturers and a kit based instead on polyA tailing followed by template switching. For our input RNA we used an equimolar VCG pool (**Methods**) with total RNA amount of either 10 or 20 pmol. All libraries were sequenced on an Illumina MiSeq using a v3 150-cycle kit. To compare our results across different kits we used the residual, a simple metric for bias, obtained by calculating per-VCG sequence counts per million (CPM) and then the per-VCG member deviation from its expected value (**Methods**). We plotted the spread of residuals for each library preparation method and observed that tinyRNA-seq showed the least spread of all methods (**Figure 2A, Table S1**), with an interquartile range (IQR) of 0.6, indicating low bias with few outliers. All commercial kits tested showed high bias (IQRs of > 2), with even the best performing kit, Supplier N (2025), having an IQR of 2.5 (**Figure 2A and Figure S1**).

**Figure 2:**
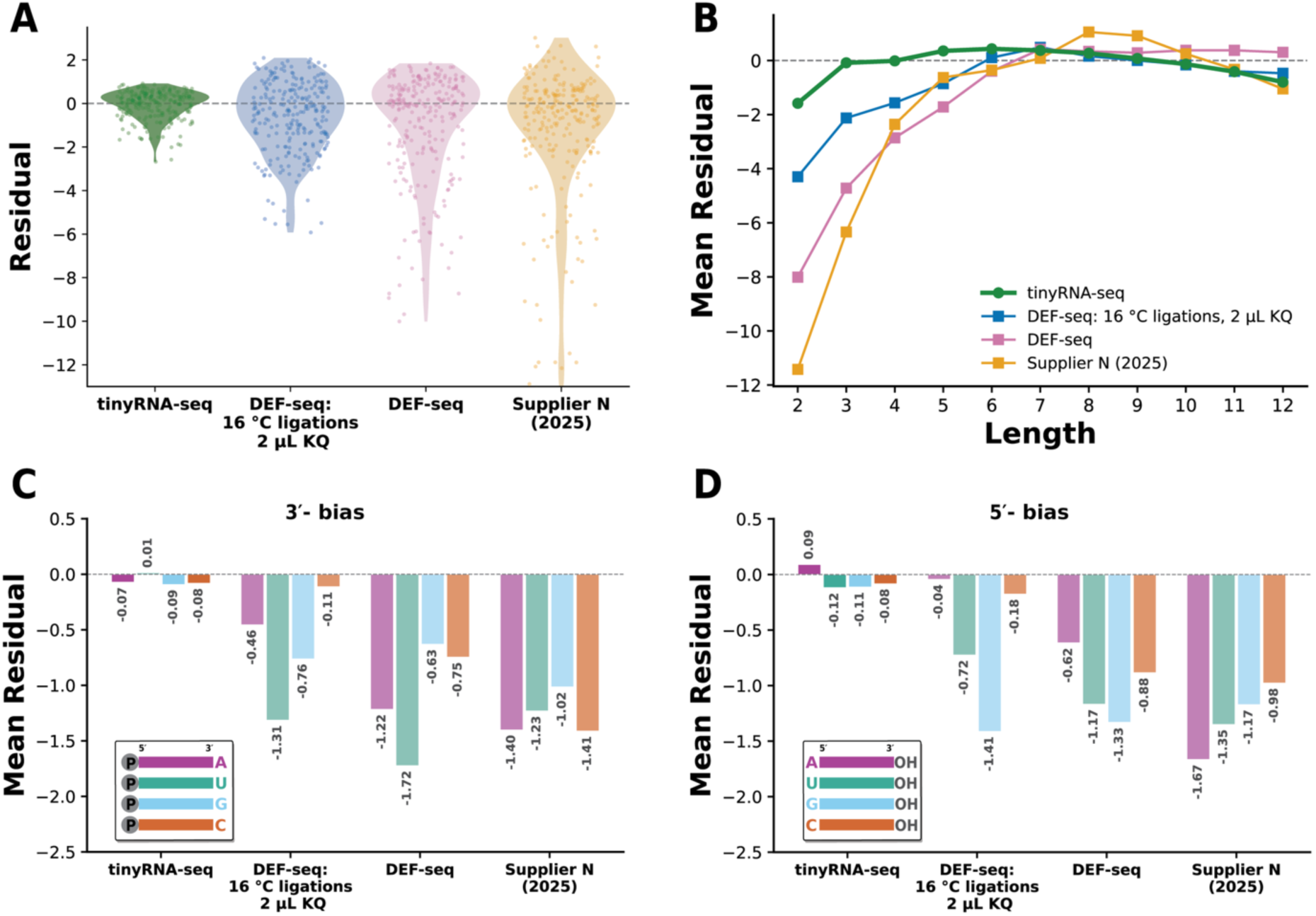
tinyRNA-seq minimizes systematic length- and primary sequence-dependent capture bias for primitive RNA genomes. Illumina sequencing libraries were prepared from virtual circular genome (VCG) tinyRNAs (a library of 247 unique 2-12 nt RNA oligos) using different ligation-based methods: tinyRNA-seq, DEF-seq (a literature-derived protocol) and a commercially available small RNA sequencing kit. (**A**) Violin plots of per-VCG species read counts for each library preparation method (colour) shown here as a residual. Residual corresponds to the deviation from expected read counts of our known VCG input pool with each integer unit corresponding to a 2-fold change from its expected read count indicated by the dashed line at 0. tinyRNA-seq shows the distribution closest to expected and therefore the least overall capture bias, DEF-seq and Supplier N (2025) show VCGs captured both above and below 0 indicating that these methods simultaneously over count and under count tinyRNAs and introduce bias. (**B**) Mean residual per VCG sequence length for each library preparation (colour). The dashed line at zero indicates unbiased recovery. Deviation as length decreases reflects length-dependent capture bias showing that methods based on defined adaptors (DEF-seq, Supplier N (2025)) fail to efficiently capture RNAs shorter than 5 nt. (**C**)–(**D**) Mean residual per-VCG sequence grouped by terminal nucleotide (A: purple, U: green, G: blue, C: orange). Bars are coloured by base identity and annotated with their residual value. (**C**) 3′-end bias is lowest for tinyRNA-seq with effectively no terminal base influencing its capture. DEF-seq shows a characteristic bias against tinyRNAs that have U termini. (**D**) 5′-end bias is also lowest overall for tinyRNA-seq.

We expected that the length of RNA input oligonucleotides was the main cause of this spread. Accordingly, we calculated the mean residual for each length of our input pool and observed that commercial kits captured RNAs shorter than 5 nt with 4-fold less recovery than expected (**Figure 2B, Figure S2**). Using even the best performing kit, Supplier N (2025), we often failed to recover many 2 nt VCG sequences.

Our comparison assay using defined adaptors (DEF-seq) also involves two sequential enzymatic ligation steps (**Figure 1 B-C**) and when we increased the amount of enzyme (2 µL T4 RNA ligase 2 KQ) and lowered incubation temperatures (16 h at 16 °C, compared to commercial kits normal incubation of 1 h at 25 °C), we also observed a length selective bias indicating that despite longer incubation times at lower temperatures, common approaches in the literature, using defined adaptors may be insufficient for unbiased capture of tinyRNAs. Since 22% of our VCG oligonucleotides are shorter than 5 nt we were pleased to observe that with our tinyRNA-seq protocol, all lengths were captured well with only minor underrepresentation of 2 nt species, with these species deviating less that 2-fold from expected (**Figure 2B, Figure S2**).

Following literature reports that primary sequence can also affect ligation efficiency, we calculated the mean residual for each 3′- and 5′-terminal base that would be involved in the first ligation and second ligation respectively (**Figure 2C-D, Figure S3-4**)[46]. In our VCG design, 3′-nucleotide identity is evenly distributed at each lengths allowing a length-agnostic investigation of ligation junction bias. When examining our results for the commercial kits we tested and DEF-seq, the mean residuals for the 3′-end base (all <0; showing bias in the ligation junction during ligation to **3pAd**) are broadly consistent with previous results on miRNA ligation junctions, indicating that ligase preference for G/C over A/U junctions is still relevant for tinyRNAs. The magnitude of bias differed with Supplier I and R exhibiting 10-fold greater bias than Supplier N (2025) and DEF-seq (**Figure 2C, Figure S3**).

Previous studies have established that the use of degenerate bases can mitigate this ligation junction specificity and we were pleased to observe that our degenerate 4 nt region in the tinyRNA-seq **3pAd** (**Figure 1B**) demonstrated 17-fold less bias for ligating tinyRNAs that have A at their 3′-end with residuals of −0.07 versus residuals of −1.22 and −1.20 for DEF-seq and Supplier N (2025), respectively (**Figure 2C**).

It is thought that T4 RNA ligase 1 has less primary sequence bias for input RNA 5′-termini but we nonetheless observed a similar decrease in bias when using our degenerate tinyRNA-seq adaptors. The tinyRNA-seq mean residual for A was 0.09, versus −0.62 and −1.45 for DEF-seq and Supplier N (2025) (**Figure 2D**). We next examined the entire VCG sequence space and noticed that tinyRNA-seq did not exhibit any systematic sequence-based bias (**Figure S5**) with almost all sequence space captured within than 2-fold than expected. Our VCG genome is comprised of a sense and antisense strand, and we noticed that both DEF-seq and commercial kits showed significant antisense strand bias which we did not observe for tinyRNA-seq (**Figure S6**). Thus tinyRNA-seq minimizes ligation bias even for tiny RNAs. This is critical as biased recovery would lead to loss of information at protogenome oligonucleotide termini.

### tinyRNA-seq performs competitively on biological RNA samples

#### Performance on biological RNA

We wondered if our tinyRNA-seq protocol would also capture small RNAs (sRNAs) from biological sources with minimal bias. There is currently great interest in the sequencing and identification of small RNAs stemming from their integral participation in eukaryotic gene regulation and defense[35–37]. Diverse biogenesis pathways produce small RNAs ranging in length from ≤17 to ∼30 nt. Given this, we envisage tinyRNA-seq to be valuable for the identification and quantification of small RNAs and in particularly tiny RNAs (<17 nt) such as cityRNAs[39, 51]. Therefore, we examined the capacity of tinyRNA-seq to capture biological small RNAs. We extracted small RNA from mouse brain tissues using the mirVana™ miRNA Isolation Kit (**Supplementary Information**) and simultaneously prepared sequencing libraries using tinyRNA-seq (with a 10x adaptor dilution to account for lower RNA input) alongside 3 commercial kits: Suppliers T, R, and N (**Supplementary Information**).

We captured 1781 distinct small RNAs including but not limited to miRNA, piRNA and snoRNA, (**Figure 3A, Table S2**). In comparison, the three commercial kits (Supplier T, R, and N) identified 1539, 1642 and 1221 unique small RNAs respectively. We then analyzed our recovered RNAs in terms of the abundance and number of small RNA classes that are known to be present in mouse brain tissues. tinyRNA-seq performed competitively with commercial kits capturing 488 unique miRNAs, compared to the commercial kits whose unique species detection ranged from 376 to 498 (**Figure 3B**). This suggests that tinyRNA-seq can reliably sequence miRNA at the same depth and sensitivity as commercial kits without significant optimization for biological RNAs.

**Figure 3:**
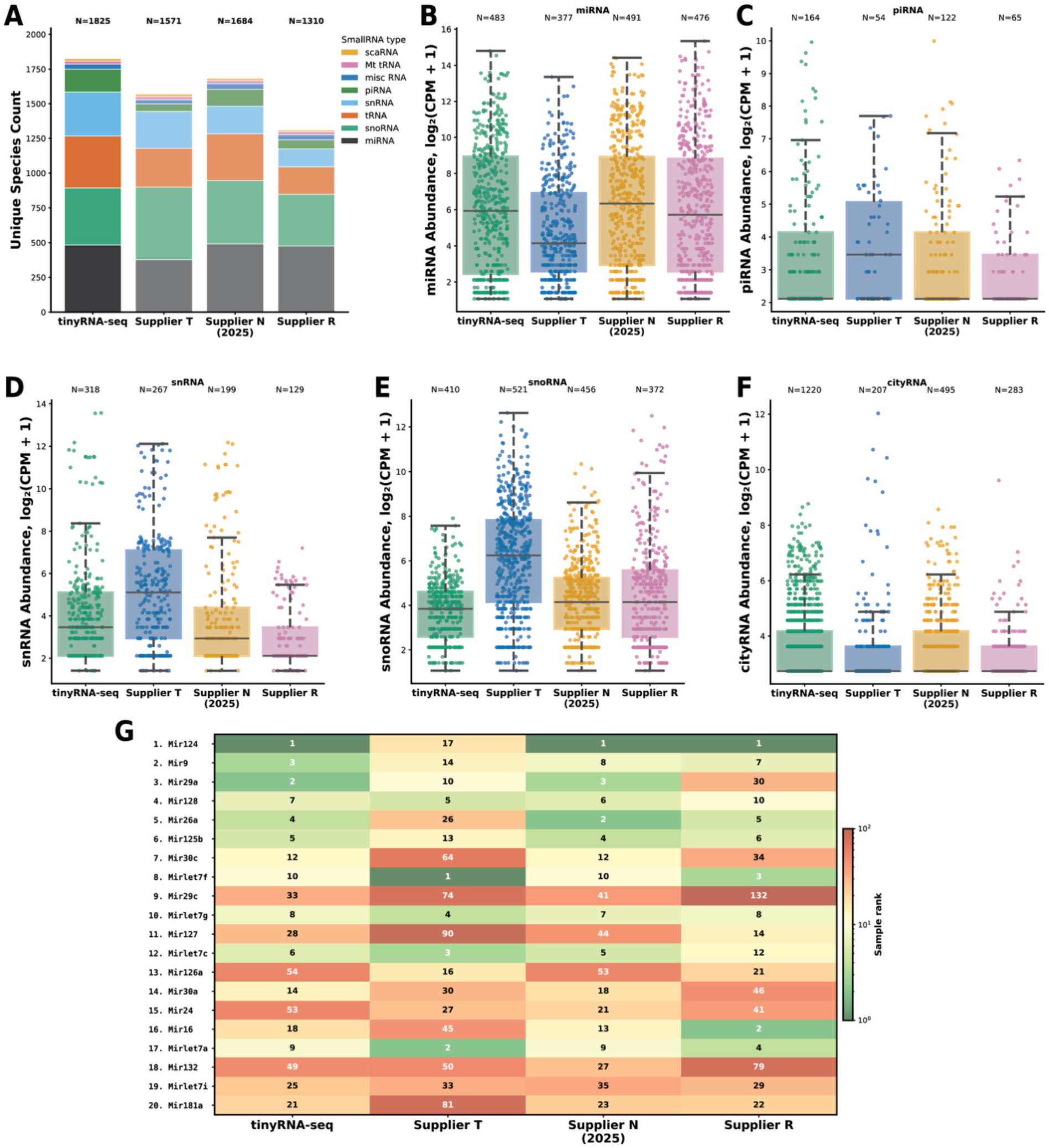
tinyRNA-seq captures biological sRNA species competitively compared to three commercial kits. Using enriched small RNAs from mouse brain tissue we tested tinyRNA-seq against 3 commercial kits designed for small RNAs. A detection threshold of greater than 1 count per million (CPM) was used for analysis. (**A**) Comparison and distribution of all unique small RNAs between tinyRNA-seq and three commercial kits: Supplier T, Supplier N (2025), and Supplier R. Stacked bar chart of total (CPM) for each method, broken down by small RNA biotype; tinyRNA-seq captures the highest number of unique small RNA species. (**B**)-(**F**) Per-species CPM distributions by small RNA biotype. Strip plots of CPM (log2 scale) for each biotype across library preparation methods. Each point is one annotated species coloured by experiment. (**B**) miRNA. (**C**) piRNA. (**D**) snRNA. (**E**) snoRNA. (**F**) cityRNA. (**G**) Heatmap of top 20 absolute abundance ranked miRNAs for all methods compared to the SmRNAQuant database, a database of quantified miRNA in specific tissues. Colour indicates sample rank on a log scale (green: correctly ranked, red: incorrectly ranked). Consistent green across a row indicates that a miRNA is faithfully recovered at high rank.

Intrigued by the possibility that tinyRNA-seq might capture low abundance species more efficiently and with less bias than commercial kits, we looked for piRNAs, spliceosomal small nuclear RNAs (snRNAs), and small nucleolar RNAs (snoRNA). piRNAs are more abundantly expressed in the germline than brain tissue and so provided a challenge to the sensitivity of our method. TinyRNA-seq identified 106 unique piRNAs whereas the commercial kits detected from 19 to 77 piRNAs (**Figure 3C**). We also identified 321 unique snRNAs using tinyRNA-seq whilst Suppliers T, R, and N identified a range of 100 to 267 unique snRNAs (**Figure 3D**). TinyRNA-seq captured 418 unique snoRNAs which is comparable to the performance of the commercial kits that detected from 388 to 519 unique snoRNAs (**Figure 3E**). Interestingly, the abundance of snRNAs and snoRNA captured using Supplier T is significantly higher than that of all other methods (**Figure 3D-E**). This is likely an artifact of Supplier T library preparation which relies on template switching reverse transcription that is reported to introduce false positive isoforms and truncated products[52, 53]. Across the four classes of small RNAs we investigated, tinyRNA-seq’s median CPM is higher or comparable to that of the commercial kits. Higher median CPM suggests the genuine capture of abundant small RNAs, above the noise and false positives that are more likely with low median CPMs. In addition to these four classes of sRNA, tinyRNA-seq also performs competitively to capture rRNA and tRNAs (**Figure S7A-B**). Recently an emerging class of small RNAs known as cleavage-inducing tiny guide RNAs (cityRNAs) has been reported[39]. CityRNAs are the 14-17 nt long products of exonuclease activity on mature miRNAs which are believed to participate in Argonaut protein (AGOs) mediated gene regulation. Their short length, within the range of detection for tinyRNA-seq, and novel biochemistry prompted us to examine our data for putative detections. We filtered for possible cityRNAs within datasets based on length of any unassigned reads (**Methods**). We report that tinyRNA-seq identified 1220 unique potential cityRNAs (≤17 nt) while commercial kits only identified between 74 to 295 potential cityRNAs (**Figure 3F**).

To test whether tinyRNA-seq was performing reliably we compared the top 20 most abundant miRNAs we captured to SmRNAQuant, an absolute quantification database[43]. We observed the ranked abundance of miRNA species captured by tinyRNA-seq and Supplier N correlates well with SmRNAQuant (**Figure 3G**, spearman rho = 0.74 vs Supplier N (2025) = 0.67), thereby suggesting that tinyRNA-seq can be reliably used for absolute small RNA quantification while Suppliers T and R underperform in terms of absolute quantification of miRNAs.

We have shown that tinyRNA-seq is compatible with biological small RNA sequencing and benchmarks competitively against commercial kits that optimized for biological samples. TinyRNA-seq was developed and intended for the quantification of synthetic short RNAs without the end and internal modifications that are common for biological sRNAs. Despite lacking enzymatic end-repair, our method detects a comparable or higher count of unique sRNA species across multiple sRNA classes including miRNA, piRNA, snRNA, and snoRNA in mouse brain tissues.

### tinyRNA-seq enables detection of *de novo* oligonucleotide generation

All current approaches to sustaining a protogenome, including our virtual circular genome model, are reliant on a supply of new primers to sustain growth. We wondered if tinyRNA-seq would allow us to study *de novo* oligonucleotide generation. Previous work has shown that RNA monomers can spontaneously polymerize without a template, especially when activated as imidazolides[54, 55]. We wondered if 2-aminoimidazolium (2AI)-bridged dinucleotides (N*N) could undergo primer-independent polymerization, thereby providing a pathway to feed a VCG-like system with new primers (**Figure 4A**). To investigate this possibility, we incubated 2AI-bridged dinucleotides (either preequilibrated mixtures of equimolar N*N or one homodimer i.e. G*G) with and without an oligonucleotide template and used tinyRNA-seq to capture and sequence newly formed small RNA fragments. When a mixture of N*Ns was incubated (16 h, RT) at different concentrations (10, 20 or 40 mM) in the absence of an oligonucleotide template we observed a distribution of *de novo* oligonucleotides from 2 to 6 nt (**Figure 4B**). Length distribution skewed longer as bridged dimer concentration increased (30% 3-mer at 10 mM, 41% 3-mer at 40 mM). G-rich products were the most abundant and even oligo-G products such as GGGG were detected (**Figure S8**) suggesting enhanced reactivity for G*G bridged dinucleotides. Even though G*G should be only 10% of the equilibrated mixture of N*N it contributed the majority of the *de novo* oligonucleotide products. When we incubated N*N (20 mM) with a homopolymer template (C_10_, 2 µM) we observed an altered product distribution with a greater proportion of longer oligo-G oligos (**Figure 4C, Figure S8A**) consistent with a template-directed mechanism. Interestingly when we incubated N*N with a G_10_ template we did not observe a commensurate increase in C-rich products (**Figure S8B**). We also wondered if we could further drive the formation of longer products by incubating only G*G on a C_10_ template (**Figure 4D**). We observed a higher proportion of 3 and 4 nt long products (**Figure 4D and S8C**) compared to using an equilibrated mixture of N*N at the same concentration. The overall length distribution was also shorter, likely because the longer polyG products are insoluble (**Figure S8C**). Interestingly, some mixed sequence oligonucleotides were observed, both in the presence of the C10 template and with no template, suggesting that these products were generated in solution and not on a template. Thus, tinyRNA-seq can be used to detect the formation of <4 nt oligonucleotides in the context of *de novo* oligonucleotide generation.

**Figure 4:**
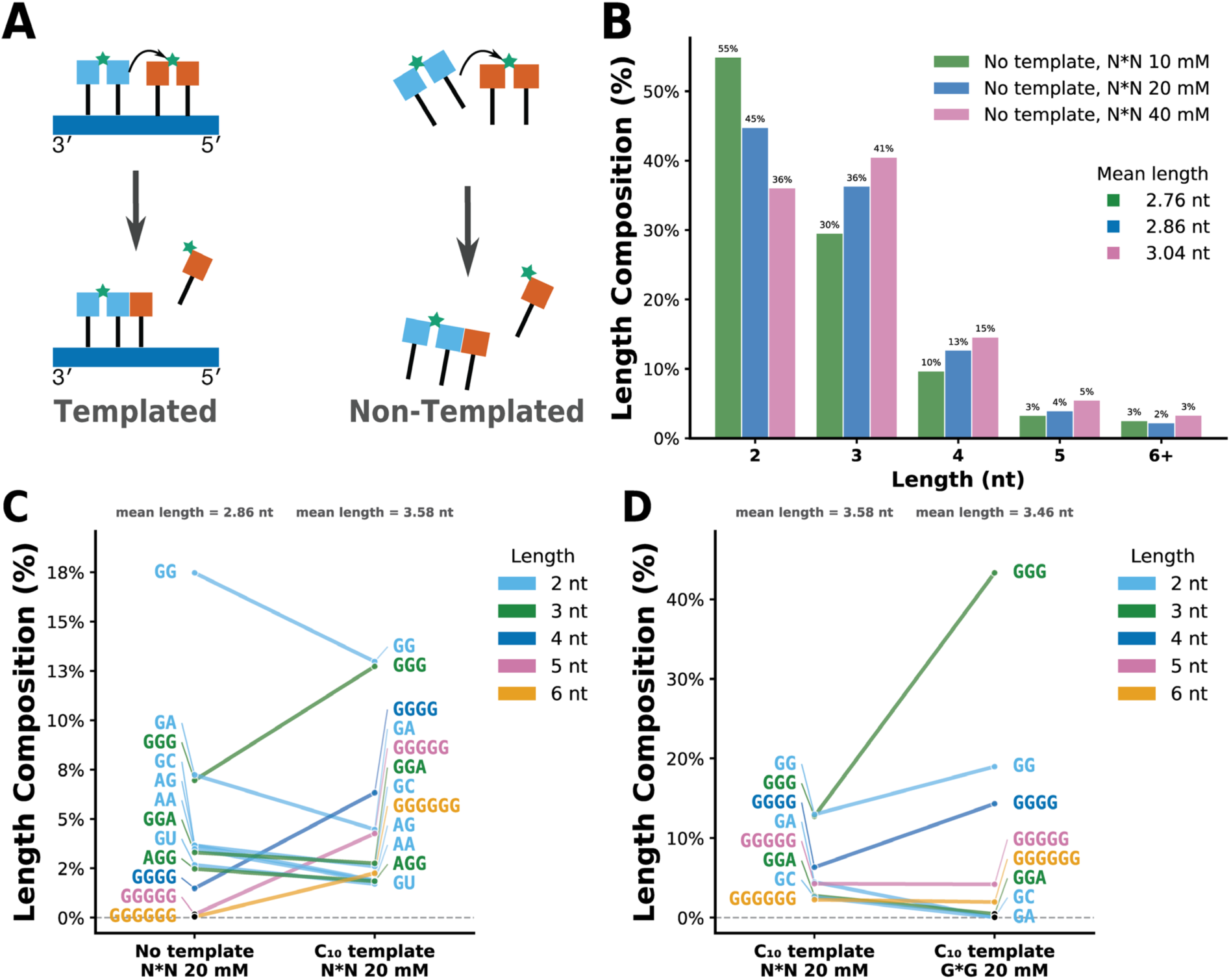
tinyRNA-seq detects de novo oligonucleotide generation and probes product distribution. tinyRNA-seq was used to sequence tinyRNA products from the incubation of reactive 5′-5′ 2-AI bridged dinucleotides (N*N) after ZipTip clean up. (**A**) Templated versus non-templated *de novo* oligonucleotide generation from N*N. (**B**) Length distribution of *de novo* oligonucleotides detected by tinyRNA-seq using N*N at 10, 20 and 40 mM. For each length (2-6 nt), bars show the fraction of total CPM contributed by sequences of that length. Increasing the concentration of N*N increases the mean product length. (**C**) Pairwise abundance comparison of individual sequence species between the no-template and C10-templated reactions at 20 mM N*N. Each line connects the same sequence across the two reactions; line position on the y-axis is the fraction of total CPM in that reaction. (**D**) Pairwise abundance comparison of C10-templated reactions with N*N versus G*G activation chemistry at 20 mM. In (**C**-**D**), sequences contributing at least 2% of total CPM in either condition are shown; lines and labels are coloured by product length, species with black markers are below experimental noise (<0.2%).

## DISCUSSION

We developed tinyRNA-seq to solve the problem of biased capture of small RNAs by current small RNA-specific sequencing pipelines. We have optimized this protocol by designing custom adaptors that simultaneously reduce ligation bias and allow for PCR duplicate collapse. Other methods, including all commercially available small RNA sequencing kits, we tested exhibited systematic length-dependent failure, with many kits failing to capture some or all of the shortest oligonucleotides (<3 nt) in our Virtual Circular Genome (VCG) test system. The sequence and distribution of these tinyRNAs is of critical interest to our exploration of the initiation and maintenance of a primitive RNA genome.

There have been many studies seeking to improve bias introduced to small RNA next generation sequencing protocols by the ligation step; tinyRNA-seq builds on this previous work but also introduces the use of degenerate regions to increase ligation junction generality as well as internal degenerate reasons that can be used for PCR deduplication. Bias introduced during PCR is becoming recognized as a larger source of bias in high-sensitivity next generation sequencing and we were keen to build in functionality to deal with this from the start as our unique molecular identifier (UMI) is installed during specific molecular ligations events, whereas UMIs are often installed during RT or PCR which has been shown to cause ambiguity during bioinformatic processing.

We explored using tinyRNA-seq for the capture of biological small RNAs due to the continued relevance of small non-coding RNAs in understanding health and disease. We were please to find that tinyRNA-seq is a method that can be adapted to the capture a diverse range of small RNA biotypes including miRNA, snoRNA piRNA and others. We found that not only was tinyRNA-seq capable of capturing a wide range of unique species it also correlated better than commercial kits with the SmRNAQuant, a tissue specific absolute quantification database of small RNAs. We expect tinyRNA-seq to be a useful tool for researchers interested in non-coding RNAs that lie at the length scale of miRNA and shorter.

An emerging class of small RNAs that are even smaller than miRNA, cityRNAs (<17 nt), currently have no good capture method to enable understanding of their distribution across tissues and species. Even the best available commercial kits struggle to capture tinyRNAs (<21 nt), like cityRNAs and our VCG genomic RNAs, which raises concerns about conclusions drawn from small RNA sequencing library preparations used to capture these tinyRNAs. tinyRNA-seq could be used by researchers interested in studying these emerging candidate non-coding RNAs.

We have previously speculated on the ability of the reactive intermediates in non-enzymatic primer extension (2-aminoimidazolium 5′-5′ bridged dinucleotides (N*N)) to polymerize spontaneously in the absence of templates to form small *de novo* oligonucleotides that can then act as primers or other components of a primitive RNA genome. tinyRNA-seq now provides us with a means of obtaining insight into both the abundance and sequence distribution of these inevitable products of prebiotic chemistry. We were interested to note that, although products of appreciable length do form in the absence of a template, the mean oligonucleotide length does increase when we incubate our N*N species with a template. Further work will be required to determine whether templated or untemplated reactions contribute more to *de novo* oligonucleotide generation in the context of VCG replication.

Finally, we have designed tinyRNA-seq, in particular our 3′-ligation adaptor (**3pAd**), to be forward compatible with nanopore sequencing and single-cell sequencing. The modular nature of our 3′-ligation adaptor (**3pAd**) design allows confidence that the most biased step of all small RNA sequencing workflows has been significantly improved and will remain significantly low-bias across future workflows. We envisage that tinyRNA-seq will be a valuable, low-bias and high-sensitivity method for researchers with diverse interests ranging from the study of primitive RNA genomes to the investigation emerging of small RNAs of clinical interest.

## ACKNOWLEDGEMENTS

We thank Alex Akhundov for the gift of imidazolium-bridged dimers, we thank the He lab at UChicago for the gift of mouse brain tissue, we thank Filip Boskovic for the gift of homopolymer templates. We thank Daniel Duzdevich and Victor Lelyveld for valuable insight and expertise. We thank Daniel Duzdevich and Filip Boskovic for comments on the manuscript, and all members of the Szostak laboratory for feedback and sharing experimental expertise. We used Claude (Anthropic; version Opus 4.8) to refine our original Python code for data processing and visualization. We reviewed all outputs and code and retain responsibility for interpreting results

## AUTHOR CONTRIBUTIONS

Ben W.F. Colville: Conceptualization [equal], Data curation [lead], Formal analysis [lead], Investigation [lead], Methodology [lead], Project administration [equal], Software [lead], Supervision [supporting], Validation [lead], Visualization [lead], Writing—original draft [lead], Writing—review & editing [equal]). Jamie Zhao: Data curation [supporting], Formal analysis [supporting], Investigation [supporting], Methodology [supporting], Validation [supporting], Writing—review & editing [supporting]). Liam Hade: Formal analysis [supporting], Software [supporting], Visualization [supporting], Writing—review & editing [supporting]). Jack W. Szostak: Conceptualization [lead], Data curation [equal], Formal analysis [equal], Funding acquisition [lead], Investigation [equal], Methodology [equal], Project administration [lead], Resources [lead], Software [supporting], Supervision [lead], Validation [equal], Visualization [supporting], Writing—original draft [equal], Writing—review & editing [lead]).

## SUPPLEMENTARY DATA

Supplementary data is available online.

