## Supplementary Information for "tinyRNA-seq: An optimized approach to sequencing tiny RNAs and primitive RNA genomes"

### Supplementary Methods

*General methods*

All chemicals were purchased from Sigma-Aldrich (St. Louis, MO) and used without purification unless otherwise noted. Phosphoramidites and reagents used for solid-phase RNA or DNA synthesis were purchased from ChemGenes (Wilmington, MA) and Glen Research (Sterling, MA). Enzymes were purchased from New England BioLabs (NEB) or ThermoFisher Scientific and used as per the manufacturer's instructions unless otherwise indicated. Reactions were performed with RNase-free water and salt solutions (Ambion) in RNase-free 1.5 ml DNA LoBind tubes (Eppendorf) or 0.2 ml PCR tubes (VWR International). We took extreme caution to prevent contamination, including the use of barrier tips for all liquid handling (Sorenson BioScience). All incubations with a specified temperature were performed in a BioRad T100 thermal cycler or Thermo Scientific QuantStudio Pro 7 qPCR machine. We used mastermixes when preparing enzymatic reactions to avoid pipetting excessively small volumes.

*Oligonucleotide synthesis*

RNA and DNA oligonucleotides were purchased from Integrated DNA Technologies (IDT, Coralville, IA) or synthesized in-house. To synthesize degenerate regions in an oligo, a premixed 3:3:2:2 molar ratio of A, C, G, and T phosphoramidites was used.

RNA Oligonucleotides were synthesized with DMT-off on a K&A H-8-SE-Oligo Synthesizer on a 1 µmol scale using pre-packed synthesis columns (Glen Research). Oligonucleotides were then cleaved from the solid support and deprotected with ammonium hydroxide solution (1.3 mL) at 65 °C for 20 mins. The mixtures were lyophilized and then for incubated with triethylamine trihydrofluoride (65 °C for 2.5 h) to remove the 2′-tert-Butyldimethylsilyl (2′-TBDMS) protecting group. The oligonucleotides were then purified by polyacrylamide gel electrophoresis (PAGE).

DNA Oligonucleotides were synthesized with DMT-off on a K&A H-8-SE-Oligo Synthesizer on a 1 µmol scale using pre-packed synthesis columns (Glen Research). Oligonucleotides were then cleaved from the solid support and deprotected with ammonium hydroxide solution (1.3 mL) at

65 °C for 20 mins. The mixtures were lyophilized and then purified by polyacrylamide gel electrophoresis (PAGE).

*Illumina MiSeq sequencing*

We typically pool 8-12 experiments when screening conditions and fewer when more than $10^6$ reads are required. Due to the lack of diversity in adaptor regions and the relatively narrow length distribution of the VCG we mix up to 30% PhiX (Illumina, FC-110-3001) into our pooled library to increase base diversity to improve sequencing quality. We used MiSeq v3 150 cycle kits (Illumina, MS-102-3001).

*Oligonucleotide adenylation*

DNA adaptors were adenylated using the 5′-adenylation kit (NEB, E2610L). DNA adaptors with a 5′-phosphate (900 pmol) were mixed with 5′ DNA Adenylation Reaction Buffer, 1mM ATP and 20 µL of Mth RNA Ligase in a final volume of 200 µL. The reaction mixture was incubated at 65 °C for 6 h and then 85 °C for 5 min to denature the ligase before spin column purification (Zymo Research) using the following modified protocol: reactions were mixed with 1600 µL 100% ethanol and 400 µL Oligo Binding Bufffer (Zymo Research) and vortexed thoroughly. This mixture was added portion wise (max 800 µL) to a Zymo-Spin™ IC Column (Zymo Research) and then either centrifuged at 12000 rcf for 1 minute or placed on a vacuum manifold, the flow through was discarded. When all the sample had been bound washing with DNA Wash Buffer was done similarly portion wise using 3200 µL of buffer. This was followed in all cases by transferring the column to a clean 1.5 mL sample tube and spinning dry for 2 mins 30 s at 12000 rcf. Finally, the now dry spin column was transferred to a new 1.5 mL sample tube and 20 µL water was added directly to the substrate before incubation for 1 min before spinning to collect the DNA which was either used immediately or frozen at -30 °C for future use.

*VCG-sequencing using commercial kits*

Sequencing our VCG pool using commercial kits as a benchmark was carried out according to all manufacturer's directions with a total RNA input of either 10 or 20 pmol VCG pool. After the PCR in each respective kit the dsDNA concentration was measured using an Agilent TapeStation (D1000 reagents), libraries were then pooled and sequenced on an Illumina MiSeq using v3 150-cycle kits.

*DEF-seq*

DEF-seq is a control protocol for small RNA sequencing. It uses a 5′-adaptor that contains an extra 4 nt at the 3′-end to prevent Illumina sequencing from starting with the input RNA.

Input RNA was first ligated to **3pAd-DEF** (**Table S3**) using T4 RNA ligase 2 (truncated, KQ) according to the manufacturer's instructions except that the ligation reaction was carried out for 20 h at 25 °C. Excess adaptor was removed using 5′-Deadenylase and RecJf and purified using a Zymo Oligo clean and concentrate kit according to manufacturer's instructions. Next **5pAd-def** (**Table S3**) was ligated using T4 RNA ligase 1, according to manufacturer's instructions and the ligation product was purified using a Zymo Oligo clean and concentrate kit. The reverse transcription primer (**RT_Pri**, **Table S3**) was annealed and reverse transcription carried out using Protoscript II according to manufacturer's instructions. The cDNA was amplified using Q5 Hot Start High Fidelity polymerase (NEB) for 16 cycles using the following program: initial denaturation at 98 °C for 30 seconds. Cycles consisted of denaturation at 98 °C for 10 seconds, annealing and extension at 62 °C for 40 seconds. A final extension step was carried out at 72 °C for 2 minutes before holding at 4 °C. We purified dsDNA libraries using preparative 3% TBE (w/v) agarose gels (UltraPure™ Agarose, Invitrogen,) and run at 90 V for 90 mins before using Freeze and Squeeze kits (BioRad) to recover our libraries. The concentration was measured using an Agilent TapeStation (D1000 reagents), libraries pooled and sequenced on an Illumina MiSeq using v3 150-cycle kits.

*Imidazolium-bridged dimers*

Solutions containing all 10 imidazolium-bridged dimers were prepared by allowing a solution containing 10 mM of each purified homodimer (A*A, U*U, G*G, C*C) to equilibrate for 2h on a rotor as previously reported[1].

*De novo oligonucleotide generation reactions*

*De novo* oligonucleotide generation reactions were carried out similarly to non-enzymatic primer extension reactions previously reported[1, 2]. Two main types of reactions were carried out: with oligonucleotide templates and without oligonucleotide templates. We used template oligonucleotides without a 5′-phosphate to prevent ligation during sequencing preparation. We also used either all 10 2-aminoimidazolium bridged dinucleotides (N*N) or an individual homodimer (e.g. G*G). In all cases reactions contained 20 mM imidazolium-bridged dinucleotide and were incubated in 200 mM Tris pH 8.0, 100 mM $MgCl_2$ and when appropriate 2 µM template oligo and allowed to react for 16 h before quenching the reaction with 200 mM EDTA and purification for sequencing.

**Post *de novo* oligonucleotide generation purification**

To remove excess salt and to concentrate *de novo* synthesized oligonucleotides samples for sequencing we utilized a modified ZipTip® procedure. Samples were processed through a sequential wash protocol using a silica-bound tip (C18 resin, bed volume 0.6 µL, tip volume 10 µL, Millipore Sigma, ZTC18S) and six wells of a PCR tube strip or plate per sample. Each well contained, in order: (i) 50 µL acetonitrile (MeCN) for tip pre-wetting, (ii) 50 µL 100 mM triethylammonium acetate (TEAA) for equilibration, (iii) 15 µL sample, (iv) 50 µL 100 mM TEAA wash, (v) 50 µL water wash, and (vi) 50 µL 1:1 MeCN:water elution buffer. For each position, the solution was aspirated and dispensed at least 20 times with a P10 pipette to ensure adequate contact between the silica and the solution, taking care not to allow the silica bed to dry between transfers. No liquid was carried over between positions; only the silica-bound material was advanced through the strip. Eluted samples were dried to completion in a Speedvac and resuspended for subsequent sequencing. This can also be done with a multichannel pipette to increase throughput but increases the difficulty of accurate pipetting to keep the silica bed wet throughout.

*De novo oligonucleotide generation sequencing*

To sequence purified products of *de novo* oligonucleotide generation we used tinyRNA-seq as described in the methods section with no modifications.

**Bioinformatics for VCG-sequencing**

For consistency, sequencing libraries prepared with the VCG pool as input raw sequencing data was processed using a custom Snakemake[3] pipeline. Each sample was analyzed independently and in parallel starting from an input .tsv file. The pipeline proceeds through five sequential steps: quality filtering, read parsing, VCG assignment, UMI deduplication, and normalization. Due to the polyA tail in libraries prepared using kits from Supplier T reads were pretrimmed using cutadapt (with -u 3 -a AAAAAAAAAA -m 2 as flags) before passing trimmed fastqs to the rest of the pipeline.

*Quality filtering*

Raw reads were first subjected to quality filtering. Per-base Phred scores were converted to error probabilities, averaged across all bases in the read, and converted back to a single mean Phred score. Reads with a mean Phred score below 30 (corresponding to a mean base-call error probability of 0.1%) were discarded.

*Read parsing and adaptor trimming*

The structure of each sequencing library depends on the preparation protocol, which determines the arrangement of the RNA insert, flanking adaptors, and any degenerate regions. These layouts are specified per-experiment in the .tsv manifest and are defined in a configuration file (adaptor_schemas.yaml). For tinyRNA-seq samples we used the read_layout of FOUR_N_MADAP, for DEF-seq samples we used DEFINED_NORMAL, for Supplier N (2025), N, I and R we used FLUSH_NORMAL with adaptor sequences from the manufacturer (see README file for an in-depth explanation). For parsing tinyRNA-seq libraries, the constant region of the

**3pAD** can be matched in each read with variable error tolerance. We typically allowed 0 errors to increase stringency but our code allows any numbers of errors to be tolerated. Using this landmark all the remaining regions of unknown sequence, but of fixed length, can be recovered from their position in the read. They are: Terminal Randomized Region (**TRR**), Binding Region (**BR**) Multiplex-Adaptor-Code (**MAC**), Internal Randomized Region (**IRR**) and Spacer (**SP**). Reads that do not conform to the expected structure (e.g. adaptor not found, insert out of expected length range) are flagged and excluded from further analysis. Two broad categories of library layout are supported: those containing degenerate regions (used as unique molecular identifiers, described below) and those without. Layouts lacking random regions skip the UMI-deduplication step.

*VCG assignment*

The RNA insert from each parsed read is compared against our VCG list (**Table S4**). Reads whose insert exactly matches a VCG in the reference list are retained; all others are excluded.

*UMI deduplication*

To correct for PCR amplification bias, reads are deduplicated using their unique molecular identifier (UMI) a short random nucleotide tag incorporated during library preparation that labels each original RNA molecule with a distinct sequence. We used an approach analgous to current literature standard UMI-tools[4]. Reads are first grouped by their VCG assignment, so only reads from the same species are ever compared. Within each group, UMI sequences are clustered using the directional method: a UMI is merged into a cluster only if its count is less than half that of the cluster root, the “2x rule”, ensuring that an abundant molecule is never incorrectly absorbed into the cluster of a different molecule. After deduplication, each cluster is represented by a single row with a count recording how many raw reads it comprised. Downstream counting uses the number of distinct UMI clusters (unique molecules), not the raw read count. Library layouts without degenerate regions bypass this step entirely and are passed through unchanged.

*Normalization*

Deduplicated read counts for each VCG were normalized to counts per million (CPM) to allow comparison across samples with different sequencing depths. The normalized output is the primary input for all downstream statistical analysis and visualization.

**Optional modules**

*Structural analysis*

Two cofold analyses were performed using the Vienna RNA 'rnacofold' package[6] to model RNA secondary structure at the ligation junction. For each read, the relevant regions flanking the ligation junctions were reconstructed from the parsed regions and submitted to rnacofold, which returns the predicted minimum free energy secondary structure in dot-bracket notation. The first ligation cofold analysis models the structural state of the RNA 3′ end in complex with the 3′-adaptor (**3pAd**). Each junction is classified into one of five structural states: open (both strands unpaired), vcg_stem (the VCG folds back on itself), adapt_stem (the adaptor forms an internal stem), capture (the VCG and adaptor base-pair with each other, forming a ligation-competent heterodimer), or complex (a mixed heterodimer and internal stem topology). The capture state is expected to favour efficient ligation by presenting free ends of both species.

The secondary structure analysis models the ligation junction between the 5′-adaptor **5pAd** and the VCG plus the **3pAd**, capturing potential structures during ligation via T4 Rnl1. The left strand is the 5′-adaptor extension concatenated with the per-read 4N degenerate region (for defined adaptor layouts any defined 5′ overhang). The right strand is the VCG sequence concatenated with 3′ adaptor as sequenced. The same 16-classes as previously described by Zhuang et al. 2012 were applied to the dot-bracket output. For each read, the size of the unpaired loop immediately flanking the ligation junction is also recorded as a measure of junction accessibility. For large experiments exceeding 500,000 reads, a per-VCG cap of 2,000 randomly sampled reads was applied before cofolding to keep computation tractable. VCGs with fewer than 2,000 reads were used in full. Structural class distributions stabilized well below this threshold, so the cap did not materially affect the summary statistics. For library layouts with no random nucleotide regions, where the left strand is constant across all reads and the right strand depends only on the VCG and 3′ adaptor sequence, each unique strand pair is folded only once.

### Biological RNA

*RNA purification*

Small RNAs were purified from mouse brain tissue generously gifted by the He lab (Department of Chemistry) at UChicago. Approximately 20 mg of brain tissue was processed using the mirVana™ miRNA Isolation Kit (Thermo Fisher, AM1560) according to the manufacturer's instructions. We opted to use this kit based on reports that it performed best when compared to 4 other commercial kits[7]. We followed the manufacturer's optional directions to enrich for small RNAs since this was our particular interest, and used 500 µL of lysis buffer for extraction and eluted the enriched small RNAs in water. We used a TapeStation high sensitivity RNA kit to measure the RNA concentration and recovered ~ 300 ng of RNA.

*Modifications to tinyRNA-seq protocol for small RNAs from biological sources*

Due to the low concentration of small RNAs present in tissue and the even smaller amount of tiny RNAs we opted to use an input amount of 5 ng per sequencing library preparation. To account for this lower input RNA amount we diluted our **3pAd** and **5pAd** adaptors by 10x as is common for commercial kits. Additionally, we opted to not carry out thermal inactivation of enzymes to prevent degradation of RNA.

### Bioinformatics for Biological RNA

Sequencing data generated from libraries prepared from mouse brains were analyzed depending on the library preparation method. For libraries prepared using tinyRNA-seq we used our Snakemake workflow described above for quality filtering, adaptor trimming and PCR duplicate collapse. For libraries prepared using kits from Supplier N (2025) or Supplier T raw reads were trimmed using cutadapt v4.9[5] using adaptor sequences provided by the manufacturer. We used the following flags to increase stringency: *–m 18* to exclude reads less than 18 nt after trimming and *-q 30* to ensure no low quality reads were retained. For Supplier T we used cutadapt with the following flags: -u 3 -a AAAAAAAAAA -m 2.

Next, all trimmed reads were aligned to the *Mus musculus* genome (mm39/GRCm39; https://www.ncbi.nlm.nih.gov/datasets/genome/GCF_000001635.27, https://hgdownload.soe.ucsc.edu/goldenPath/mm39/bigZips/mm39.fa.gz) with Bowtie v1.3.1[8], reporting a single best-stratum alignment per read (--best --strata -k 1), permitting up to one mismatch (-v 1), and discarding reads mapping to more than 1000 loci (-m 1000). Alignments were converted to BAM, sorted and indexed with SAMtools v1.21[9] for downstream use.

Sequence counts for small RNAs were generated using featureCounts (v2.1.1, Subread package)[10] against a merged genomic annotation combining GENCODE (miRNA, snoRNA, snRNA, scaRNA, rRNA, Mt-rRNA and Mt-tRNA, and other ncRNA loci), GtRNAdb (nuclear tRNAs) and RepeatMasker (repeat-derived classes e.g. SINE, LINE, LTR).

piRNAs, which lack GENCODE loci, were counted separately by conservatively matching (no mismatches allowed, no fractional counting) genome-aligned reads unassigned by featureCounts to reference piRNA sequences (piRNAdb mouse release v1.7.6, piRNAdb.mmu.v1_7_6, together with the piRNAdb mouse "gold" set, mmu.gold) by exact full-length identity.

cityRNAs were quantified separately: trimmed reads shorter than the 18 nt minimum alignment length were set aside before Bowtie alignment and those falling within 12–15 nt were matched by exact, gapless substring identity (0 mismatches) to mature mouse miRNA sequences (miRBase-derived), taking the highest-expression isoform as the canonical sequence per miRNA; each qualifying short read was recorded against every miRNA it matched.

Aligned reads for small RNAs (miRNAs, snoRNAs etc), piRNA, and cityRNAs were randomly subsampled (fixed seed) to a common depth to enable comparison and counts were normalized to counts per million (CPM) as above to allow comparison between experiments. For all analysis we set a threshold of CPM >1 as a detection limit.

**Supplementary Figures**

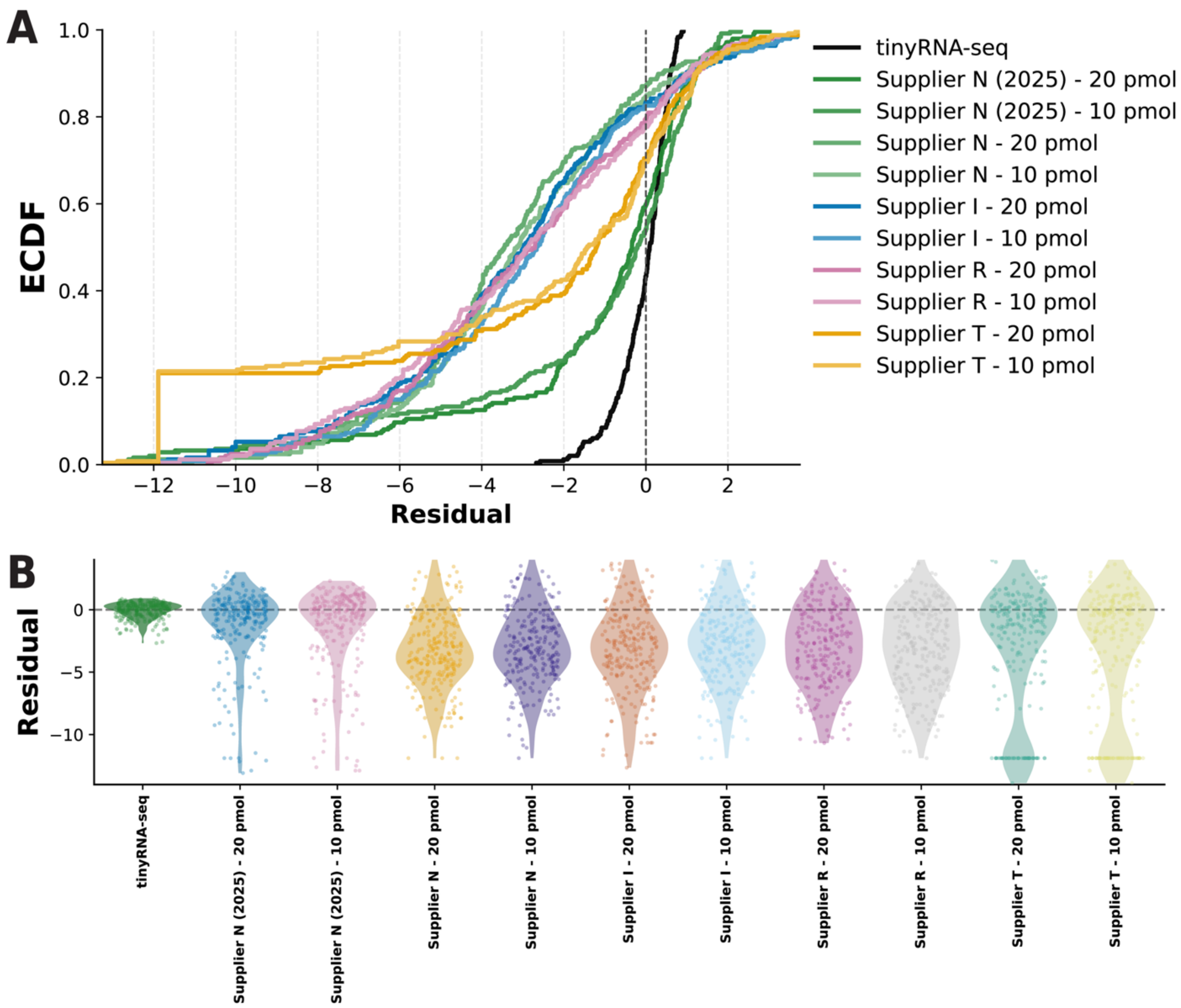


**Figure S1:** (**A**) *Per-VCG residuals for all commercial library preparation kits compared to tinyRNA-seq*. Empirical Cumulative Distribution Function (ECDF) of residuals for each sequencing run, coloured by kit supplier and input amount (10 or 20 pmol). Residual = $\log_2(\text{CPM\_observed} + 1) - \log_2(\text{CPM\_expected} + 1)$, with CPM_expected (counts per million) derived from the known equimolar VCG pool composition; each integer unit corresponds to a 2-fold change from expected. Curves centered near zero indicate accurate recovery of the VCG pool; shifts above 0 and below 0 reflect systematic over- or under-representation respectively. (**B**) Violin plots of per-VCG residuals for tinyRNA-seq compared to all commercial kits tested.

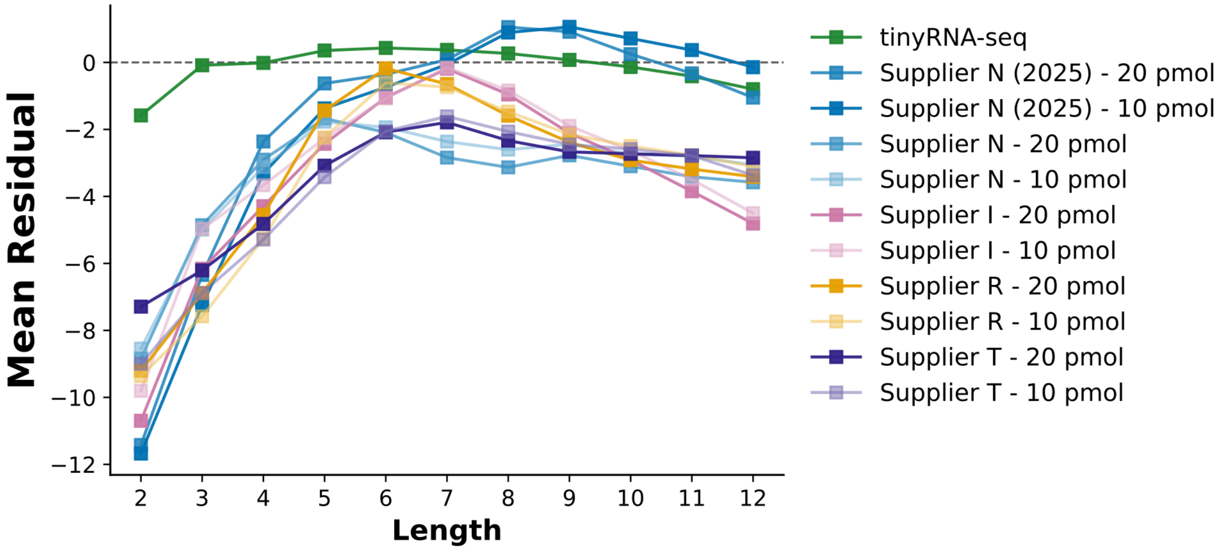


**Figure S2:** *Mean residual by VCG length across commercial library preparation kits compared to tinyRNA-seq*. Mean residual per VCG sequence length for each run, grouped by supplier (colour) with input RNA amount (20 or 10 pmol) distinguished by marker shading. The dashed line at zero indicates unbiased recovery. Systematic deviation with length reflects length-dependent capture bias.

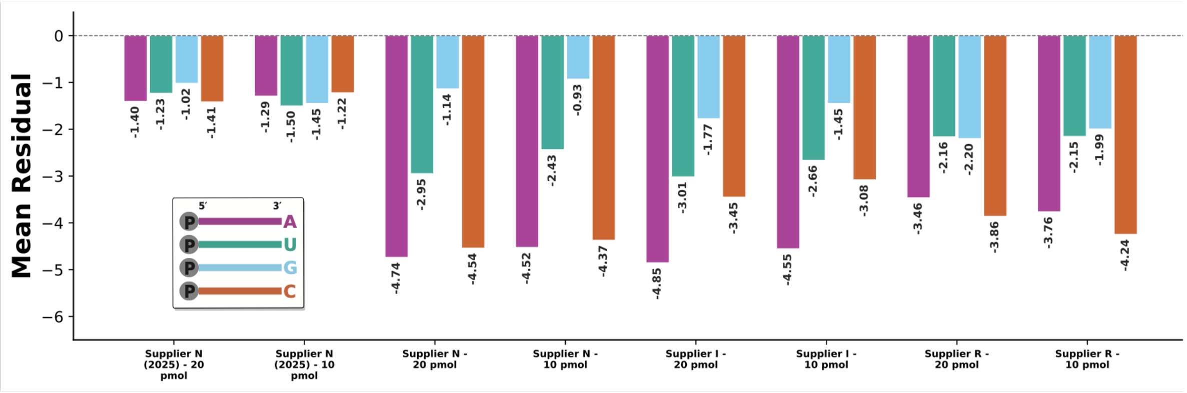


**Figure S3:** *3′-terminal nucleotide bias across commercial library preparation kits.* Mean residual per VCG sequence grouped by 3′-terminal nucleotide (A: purple, U: green, G: blue, C: orange) for each run. Bars are coloured by base identity and annotated with their residual value. Deviation from zero indicates ligation bias associated with the 3′-terminal base. Supplier T was here omitted due to end uncertainty introduced by polyA tailing.

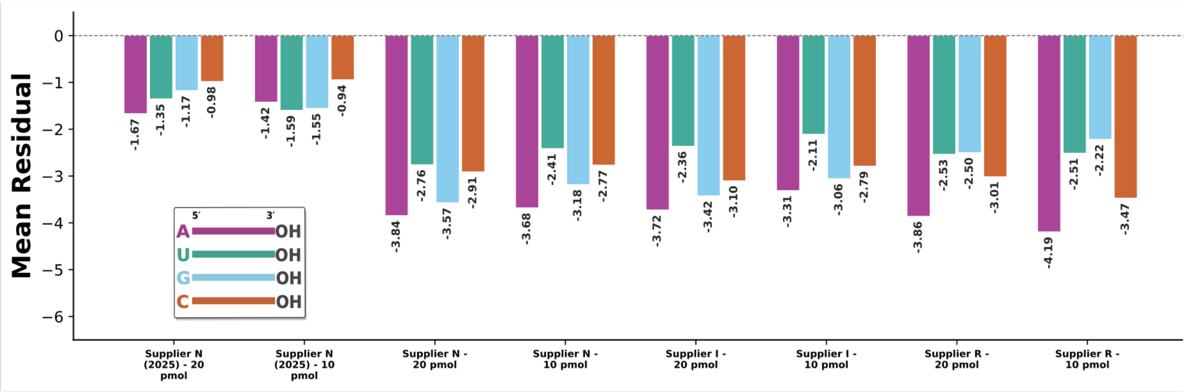


**Figure S4:** *5′-terminal nucleotide bias across commercial library preparation kits.* Mean residual per VCG sequence grouped by 5′-terminal nucleotide (A: purple, U: green, G: blue, C: orange) for each run. Bars are coloured by base identity and annotated with their residual value. Deviation from zero indicates ligation bias associated with the 5′-terminal base. Supplier T was here omitted due to end uncertainty introduced by template switching.

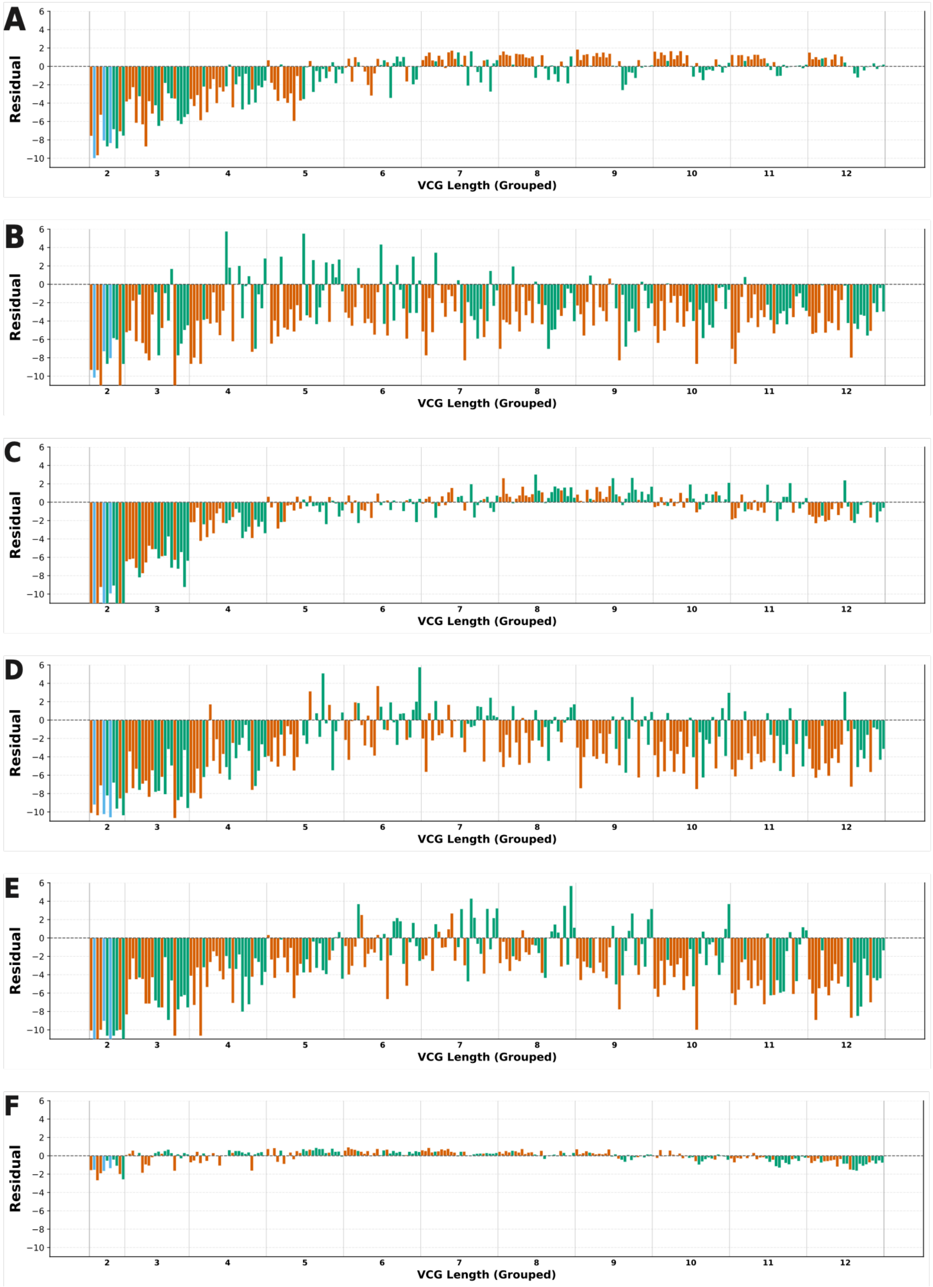


**Figure S5:** *Per-VCG residual profiles across commercial library preparation kits.* Residual for each VCG sequence shown as a bar chart for each run, coloured by strand assignment (sense: green, antisense: orange, shared: blue). Sequences are ordered and grouped by length; vertical lines demarcate lengths. Consistent positive or negative residuals within a length group indicate length-dependent capture bias. (**A**) DEF-seq. (**B**) Supplier N. (**C**) Supplier N (2025). (**D**)Supplier R. (**E**) Supplier I. (**F**) tinyRNA-seq.

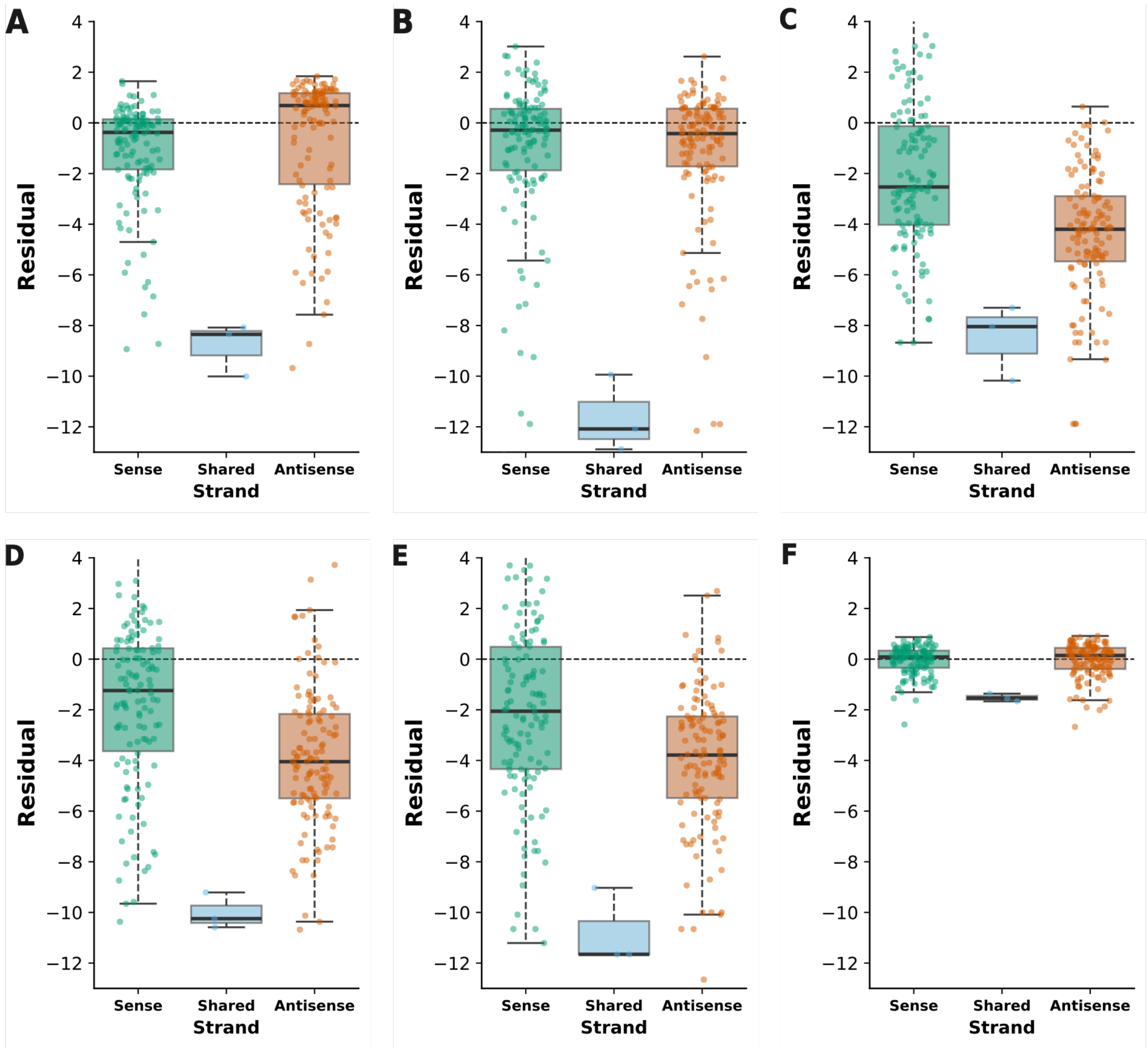


**Figure S6:** *Residual distribution by VCG strand class across library preparation methods.* Boxplots of residual for each strand class (sense: green, antisense: orange, shared: blue) shown per experiment. Horizontal lines show medians; annotated values indicate median residual per class. Systematic differences between strand classes indicate strand-dependent ligation or capture bias independent of sequence identity. (**A**) DEF-Seq. (**B**) Supplier N (2025). (**C**) Supplier N. (**D**) Supplier R. (**E**) Supplier I. (**F**) tinyRNA-seq.

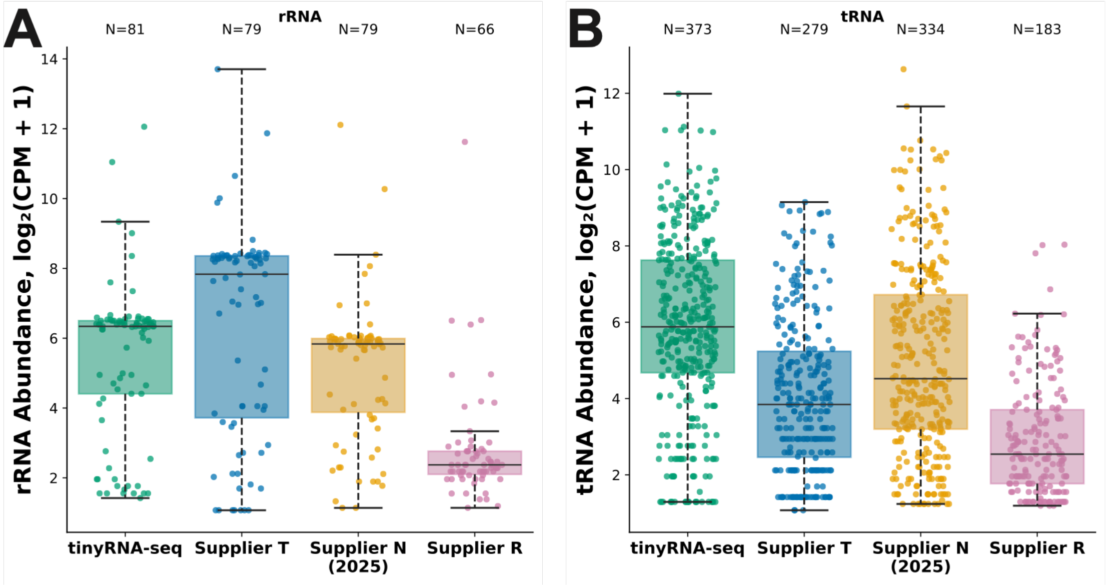


**Figure S7:** *rRNA and tRNA read distributions across small RNA libraries.* Strip plots showing CPM distributions of individual rRNA and tRNA species detected in mouse brain tissue by each library preparation. Each point represents one annotated gene/species; horizontal bars indicate the median. CPM values are plotted on a log2 scale. (**A**) rRNA. (**B**) tRNA.

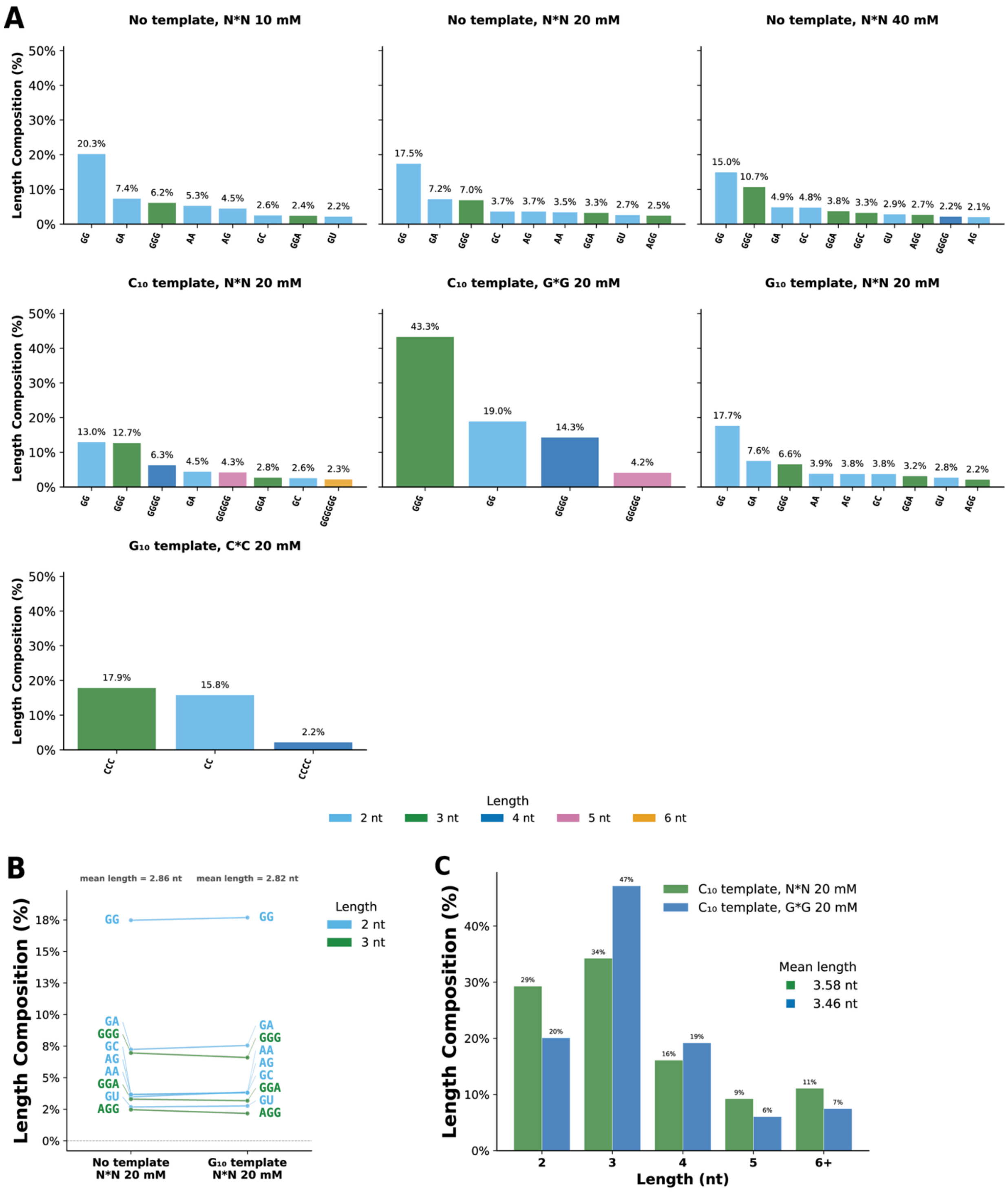


**Figure S8:** *Product distribution de novo oligonucleotide generation* (**A**) Per-experiment high-frequency products. For each reaction, all sequences contributing at least 2% of total CPM are shown, ordered by descending abundance and annotated with their CPM fraction. Bar colour denotes product length. (**B**) Pairwise abundance comparison of individual sequence species between the no-template and $G_{10}$ templated reactions at 20 mM N*N. Each line connects the same sequence across the two reactions; line position on the y-axis is the species' fraction of total CPM in that reaction. Sequences contributing at least 2% of CPM in either reaction are shown; lines and labels are coloured by product length. (**C**) Length composition of the $C_{10}$-templated reactions with N*N versus G*G activation chemistry at 20 mM. For each product length (2-6 nt), grouped bars show the fraction of that reaction's total CPM contributed by sequences of that length.

**Supplementary Tables**

| Entry | Library Preparation | VCG Species Detected | Mean Residual | Median Residual | Q25 | Q75 | IQR |
|---|---|---|---|---|---|---|---|
| 1 | tinyRNA-seq | 247 | -0.063 | 0.088 | -0.393 | 0.389 | 0.782 |
| 2 | DEF-seq: 16 C ligations, 2 µL KQ | 247 | -0.625 | -0.366 | -1.569 | 0.648 | 2.216 |
| 3 | DEF-seq | 247 | -1.018 | -0.064 | -2.074 | 0.751 | 2.825 |
| 4 | Supplier N (2025) | 247 | -1.258 | -0.341 | -1.908 | 0.562 | 2.47 |

**Table S1:** *Summary statistics for per-VCG residual distributions across library preparation methods (companion to Fig 2A)*. Residuals are defined as $\log_2$(CPM_observed + 1) − $\log_2$(CPM_expected + 1), where CPM_expected is derived from the equimolar VCG pool composition; each integer unit corresponds to a 2-fold change from expected. A residual of 0 indicates a VCG recovered at its expected abundance; negative values indicate under-representation, positive values over-representation. Values reported per experiment: n: number of VCG species contributing to the distribution, mean: arithmetic mean residual, median: 50th percentile, Q25 and Q75: 25th and 75th percentiles, IQR: interquartile range (Q75 − Q25).

| Entry | Library Preparation | miRNA | snoRNA | tRNA | snRNA | piRNA | rRNA | miscRNA | Mt tRNA | scaRNA | Mt rRNA | Total |
|---|---|---|---|---|---|---|---|---|---|---|---|---|
| 1 | tinyRNA-seq | 488 | 418 | 373 | 321 | 106 | 81 | 35 | 22 | 18 | 2 | 1864 |
| 2 | Supplier T | 376 | 519 | 279 | 267 | 26 | 79 | 30 | 22 | 20 | 2 | 1620 |
| 3 | Supplier N (2025) | 498 | 474 | 334 | 189 | 77 | 79 | 32 | 22 | 16 | 2 | 1723 |
| 4 | Supplier R | 466 | 388 | 183 | 100 | 19 | 66 | 29 | 22 | 14 | 2 | 1289 |

**Table S2:** *Unique RNA species detected per biotype across library preparation methods (companion to Fig 4A)*. Values are counts of distinct annotated RNA species detected in each experiment, grouped by Ensembl gene_type classification. Detection was defined as CPM ≥ 1 (see Methods). Biotypes shown include small RNAs: miRNA (canonical miRNAs), tRNA and mt tRNA (nuclear- and mitochondrial-encoded transfer RNAs), snoRNA (small nucleolar RNAs), snRNA (small nuclear RNAs), scaRNA (small Cajal-body-associated RNAs), and miscRNA (miscellaneous non-coding RNAs not falling into the above classes). We also report other RNAs such as rRNA (ribosomal RNA) and Mt rRNA (mitochondrial-encoded ribosomal RNAs.

| Oligo Name | Type | Source | Sequence (5′-3′) |
|---|---|---|---|
| 3pAd-CAA | DNA | In House | /5rApp/NNNNAGATCGCAANNNNCTAGATCGGAAGAGCACACGTCTAAdUGAAT/3ddC/ |
| 3pAd-GTT | DNA | In House | /5rApp/NNNNAGATCGGTTNNNNCTAGATCGGAAGAGCACACGTCTAAdUGAAT/3ddC/ |
| 3pAd-DEF | DNA | In house | /5rApp/AGATCGGAAGAGCACACGTCT/3ddC/ |
| 5pAd-4N | RNA | In House | GUUCAGAGUUCUACAGUCCGACGAUCNNNN |
| 5pAd-DEF | RNA | In House | GUUCAGAGUUCUACAGUCCGACGAUCUCUA |
| RT_Primer | DNA | IDT | AGACGTGTGCTCTTCCGATCT |
| PCR Primer 1 (SR Primer) | DNA | NEB/IDT | AATGATACGGCGACCACCGAGATCTACACGTTCAGAGTTCTACAGTCCG**-s-**A |
| PCR Primer 2 (Index Primer) | DNA | NEB | CAAGCAGAAGACGGCATACGAGATNNNNNNGTGACTGGAGTTCAGACGTGTGCTCTTCCGATC**-s-**T |
| $C_{10}$-Template | RNA | In House | FAM-CCCCCCCCCC |
| $G_{10}$-Template | RNA | In House | FAM-GGGGGGGGGG |

**Table S3:** *Table of oligonucleotide sequences used in this study*. All termini free OH unless otherwise noted.

Abbreviations:

| | |
|---|---|
| N | Degenerate position |
| 5rApp | 5′-adenylation |
| dU | deoxyuridine |
| 3ddC | 3′-dideoxycytidine |
| -s- | phosphorthioate linkage |
| FAM | 5′-fluorescein |

| | | | | | | | | | |
|---|---|---|---|---|---|---|---|---|---|
| v201 | pAC | v301 | pACA | v401 | pACAC | v501 | pACACG | v601 | pACACGC |
| v202 | pAU | v302 | pACC | v402 | pACCA | v502 | pACCAC | v602 | pACCACA |
| v203 | pCA | v303 | pACG | v403 | pACGC | v503 | pACGCA | v603 | pACGCAU |
| v204 | pCC | v304 | pAUC | v404 | pAUCA | v504 | pAUCAC | v604 | pAUCACC |
| v205 | pCG | v305 | pAUG | v405 | pAUGC | v505 | pAUGCG | v605 | pAUGCGU |
| v206 | pGA | v306 | pCAC | v406 | pCACA | v506 | pCACAC | v606 | pCACACG |
| v207 | pGC | v307 | pCAU | v407 | pCACC | v507 | pCACCA | v607 | pCACCAC |
| v208 | pGG | v308 | pCCA | v408 | pCACG | v508 | pCACGC | v608 | pCACGCA |
| v209 | pGU | v309 | pCCG | v409 | pCAUC | v509 | pCAUCA | v609 | pCAUCAC |
| v210 | pUC | v310 | pCGU | v410 | pCCAC | v510 | pCCACA | v610 | pCCACAC |
| v211 | pUG | v311 | pGAU | v411 | pCGCA | v511 | pCGCAU | v611 | pCGCAUC |
| | | v312 | pGCA | v412 | pCGUG | v512 | pCGUGU | v612 | pCGUGUG |
| | | v313 | pGCG | v413 | pGAUG | v513 | pGAUGC | v613 | pGAUGCG |
| | | v314 | pGGU | v414 | pGCAU | v514 | pGCAUC | v614 | pGCAUCA |
| | | v315 | pGUG | v415 | pGCGU | v515 | pGCGUG | v615 | pGCGUGU |
| | | v316 | pUCA | v416 | pGGUG | v516 | pGGUGA | v616 | pGGUGAU |
| | | v317 | pUGA | v417 | pGUGA | v517 | pGUGAU | v617 | pGUGAUG |
| | | v318 | pUGC | v418 | pGUGG | v518 | pGUGGU | v618 | pGUGGUG |
| | | v319 | pUGU | v419 | pGUGU | v519 | pGUGUA | v619 | pGUGUAU |
| | | v320 | pUGG | v420 | pUCAC | v520 | pUCACC | v620 | pUCACCA |
| | | | | v421 | pUGAU | v521 | pUGAUG | v621 | pUGAUGC |
| | | | | v422 | pUGCG | v522 | pUGCGU | v622 | pUGCGUG |
| | | | | v423 | pUGGU | v523 | pUGGUG | v623 | pUGGUGA |
| | | | | v424 | pUGUG | v524 | pUGUGG | v624 | pUGUGGU |

**Table S4:** *Sequences of all VCG oligonucleotides, and their associated identifying code, used in this study.* All species are 5′-phosphorylated with 3′-hydroxyls. These oligonucleotides were used in prior work by our group and were prepared by solid phase oligonucleotide synthesis in house or by IDT[1].

| | | | | | |
|---|---|---|---|---|---|
| v701 | pACACGCA | v801 | pACACGCAU | v901 | pACACGCAUC |
| v702 | pACCACCA | v802 | pACCACCAC | v902 | pACCACCACA |
| v703 | pACGCAUC | v803 | pACGCAUCA | v903 | pACGCAUCAC |
| v704 | pAUCACCA | v804 | pAUCACCAC | v904 | pAUCACCACA |
| v705 | pAUGCGUG | v805 | pAUGCGUGU | v905 | pAUGCGUGUG |
| v706 | pCACACGC | v806 | pCACACGCA | v906 | pCACACGCAU |
| v707 | pCACCACA | v807 | pCACCACAC | v907 | pCACCACACG |
| v708 | pCACGCAU | v808 | pCACGCAUC | v908 | pCACGCAUCA |
| v709 | pCAUCACC | v809 | pCAUCACCA | v909 | pCAUCACCAC |
| v710 | pCCACACG | v810 | pCCACACGC | v910 | pCCACACGCA |
| v711 | pCGCAUCA | v811 | pCGCAUCAC | v911 | pCGCAUCACC |
| v712 | pCGUGUGG | v812 | pCGUGUGGU | v912 | pCGUGUGGUG |
| v713 | pGAUGCGU | v813 | pGAUGCGUG | v913 | pGAUGCGUGU |
| v714 | pGCAUCAC | v814 | pGCAUCACC | v914 | pGCAUCACCA |
| v715 | pGCGUGUG | v815 | pGCGUGUGG | v915 | pGCGUGUGGU |
| v716 | pGGUGAUG | v816 | pGGUGAUGC | v916 | pGGUGAUGCG |
| v717 | pGUGAUGC | v817 | pGUGAUGCG | v917 | pGUGAUGCGU |
| v718 | pGUGGUGA | v818 | pGUGGUGAU | v918 | pGUGGUGAUG |
| v719 | pGUGUGGU | v819 | pGUGUGGUG | v919 | pGUGUGGUGA |
| v720 | pUCACCAC | v820 | pUCACCACA | v920 | pUCACCACAC |
| v721 | pUGAUGCG | v821 | pUGAUGCGU | v921 | pUGAUGCGUG |
| v722 | pUGCGUGU | v822 | pUGCGUGUG | v922 | pUGCGUGUGG |
| v723 | pUGGUGAU | v823 | pUGGUGAUG | v923 | pUGGUGAUGC |
| v724 | pUGUGGUG | v824 | pUGUGGUGA | v924 | pUGUGGUGAU |

**Table S4:** Continued.

| | | | | | |
|---|---|---|---|---|---|
| v1001 | pACACGCAUCA | v1101 | pACACGCAUCAC | v1201 | pACACGCAUCACC |
| v1002 | pACCACCACAC | v1102 | pACCACCACACG | v1202 | pACCACCACACGC |
| v1003 | pACGCAUCACC | v1103 | pACGCAUCACCA | v1203 | pACGCAUCACCAC |
| v1004 | pAUCACCACAC | v1104 | pAUCACCACACG | v1204 | pAUCACCACACGC |
| v1005 | pAUGCGUGUGG | v1105 | pAUGCGUGUGGU | v1205 | pAUGCGUGUGGUG |
| v1006 | pCACACGCAUC | v1106 | pCACACGCAUCA | v1206 | pCACACGCAUCAC |
| v1007 | pCACCACACGC | v1107 | pCACCACACGCA | v1207 | pCACCACACGCAU |
| v1008 | pCACGCAUCAC | v1108 | pCACGCAUCACC | v1208 | pCACGCAUCACCA |
| v1009 | pCAUCACCACA | v1109 | pCAUCACCACAC | v1209 | pCAUCACCACACG |
| v1010 | pCCACACGCAU | v1110 | pCCACACGCAUC | v1210 | pCCACACGCAUCA |
| v1011 | pCGCAUCACCA | v1111 | pCGCAUCACCAC | v1211 | pCGCAUCACCACA |
| v1012 | pCGUGUGGUGA | v1112 | pCGUGUGGUGAU | v1212 | pCGUGUGGUGAUG |
| v1013 | pGAUGCGUGUG | v1113 | pGAUGCGUGUGG | v1213 | pGAUGCGUGUGGU |
| v1014 | pGCAUCACCAC | v1114 | pGCAUCACCACA | v1214 | pGCAUCACCACAC |
| v1015 | pGCGUGUGGUG | v1115 | pGCGUGUGGUGA | v1215 | pGCGUGUGGUGAU |
| v1016 | pGGUGAUGCGU | v1116 | pGGUGAUGCGUG | v1216 | pGGUGAUGCGUGU |
| v1017 | pGUGAUGCGUG | v1117 | pGUGAUGCGUGU | v1217 | pGUGAUGCGUGUG |
| v1018 | pGUGGUGAUGC | v1118 | pGUGGUGAUGCG | v1218 | pGUGGUGAUGCGU |
| v1019 | pGUGUGGUGAU | v1119 | pGUGUGGUGAUG | v1219 | pGUGUGGUGAUGC |
| v1020 | pUCACCACACG | v1120 | pUCACCACACGC | v1220 | pUCACCACACGCA |
| v1021 | pUGAUGCGUGU | v1121 | pUGAUGCGUGUG | v1221 | pUGAUGCGUGUGG |
| v1022 | pUGCGUGUGGU | v1122 | pUGCGUGUGGUG | v1222 | pUGCGUGUGGUGA |
| v1023 | pUGGUGAUGCG | v1123 | pUGGUGAUGCGU | v1223 | pUGGUGAUGCGUG |
| v1024 | pUGUGGUGAUG | v1124 | pUGUGGUGAUGC | v1224 | pUGUGGUGAUGCG |

**Table S4:** Continued.
